# Time-resolved single-cell and spatial gene regulatory atlas of plants under pathogen attack

**DOI:** 10.1101/2023.04.10.536170

**Authors:** Tatsuya Nobori, Alexander Monell, Travis A. Lee, Jingtian Zhou, Joseph Nery, Joseph R. Ecker

## Abstract

Plant leaf intercellular space provides a nutrient-rich and heterogeneous niche for microbes that critically impacts plant health. However, how individual plant cells respond to heterogeneous microbial colonization remains largely elusive. Here, by time-resolved simultaneous single-cell transcriptome and epigenome profiling of plants (*Arabidopsis thaliana*) infected by virulent and avirulent bacterial pathogens (*Pseudomonas syringae*), we present cell atlases with gene regulatory logic involving transcription factors, putative *cis*-regulatory elements, and target genes associated with disease and immunity. We also identify previously uncharacterized cell populations with distinct immune gene expression within major developmental cell types. Furthermore, we employ time-resolved spatial transcriptomics to reveal spatial heterogeneity of plant immune responses linked to pathogen distribution. Integrating our single-cell multiomics and spatial omics data enables spatiotemporal mapping of defense gene regulatory logic with pathogen cells. Our study provides a molecularly-defined spatiotemporal map of plant-microbe interaction at the single-cell resolution.

## Introduction

Interactions between hosts and microbes can be heterogeneous for multiple reasons: (1) multicellular host tissues are comprised of diverse cell types that have distinct capacities to respond to microbes, (2) microbes can occupy niches heterogeneously distributed in the host, and (3) individual interaction events between cells may occur in an asynchronous manner in a tissue. Resolving such heterogeneity is important for understanding cellular interactions and gene regulatory mechanisms in host and microbial cells. However, the degree of heterogeneity and the contributions of the above-mentioned factors still need to be clarified.

Plant-pathogen interactions have been studied to understand molecular mechanisms underlying host immunity and pathogen virulence (Ngou et al., 2022). Plants do not have mobile immune cells or an adaptive immune system, but individual cells have the potential to mount an innate immune response upon pathogen invasion. The first layer of plant innate immunity is pattern-triggered immunity (PTI), which is activated by cell surface-localized immune receptors that recognize pathogen-derived molecules (Couto and Zipfel, 2016).

Adapted pathogens have evolved a suite of small molecules called effectors that are injected into plant cells and interfere with PTI pathways to cause diseases (Büttner and He, 2009). The second layer of plant innate immunity is a counter-defense to such effectors triggered by nucleotide-binding domain and leucine-rich repeat (NLR) immune receptors that recognize specific effectors; this immunity is called effector-triggered immunity (ETI) (Cui et al., 2015). Activation of PTI and ETI leads to massive reprogramming of gene expression regulated by many transcription factors (TFs) and co-regulators (Tsuda and Somssich, 2015). Gene regulation by TFs is encoded by *cis*-regulatory elements (CREs), DNA regions that contain clusters of TF binding sites (Marand et al., 2017). A previous study employed assay for transposase-accessible chromatin followed by sequencing (ATAC-seq) to identify accessible chromatin regions (ACRs) upon activation of immunity in bulk tissues, characterizing putative CREs involved in plant immunity (Ding et al., 2021).

Despite our understanding of transcriptional reprogramming and the underlying regulatory mechanisms during pathogen infection at the tissue scale, it is largely unknown how diverse and heterogeneous immune cell states and underlying mechanisms are at the single-cell level. Single-cell RNA-seq (scRNA-seq) has recently been applied to tackle this problem (Zhu et al., 2022; Tang et al., 2023). scRNA-seq analysis of *A. thaliana* leaf protoplasts infected by the virulent *P. syringae* pv*. tomato* DC3000 (hereafter DC3000), a model bacterial pathogen that proliferates in the intercellular space (apoplast), revealed heterogeneous expression of genes involved in immunity and susceptibility at a single time point, which could be explained by heterogeneous distribution of pathogen cells (Zhu et al., 2022). The authors reported that the expression of nearly a third of the transcriptome (7,548 genes) could be affected by protoplast generation, complicating the analysis of pathogen-inducible genes (Zhu et al., 2022). Single-nucleus omics approaches can mitigate this problem and is an alternative to protoplast-based approaches (Cole et al., 2021). Recently, single-nucleus ATAC-seq was applied to plant tissues and showed that active CREs are heterogeneously distributed across plant tissues; many of them were cell-type-specific (Marand et al., 2021). However, how pathogen infection impacts the CRE landscape at the single-cell level remains an important open question.

Linking gene expression and chromatin accessibility in single cells has the potential to reveal diverse gene regulatory logic in complex tissues. However, integration between separate scRNA-seq and scATAC-seq datasets is challenging unless well-annotated atlases of cell populations for each modality are available or strong associations between two modalities (e.g., gene body accessibility and gene expression) for each gene are known (Argelaguet et al., 2021). Although many computational methods have been actively developed to solve this problem, it can be experimentally bypassed by single-cell multiomics assays, which allow the profiling of multiple molecular modalities from the same cell (Vandereyken et al., 2023). Matched multimodal assays can serve as gold standard datasets for integrating separate unimodal datasets.

While such high throughput single-cell multiomics can define cell populations with molecular details, spatial information of cells is lost during the necessary cell (or nuclei) dissociation process, which fundamentally limits our understanding of tissue heterogeneity. A previous study that used transgenic reporter plants and live imaging showed distinct spatial expression patterns of genes involved in different defense pathways upon pathogen infection (Betsuyaku et al., 2018). Zhu et al. also employed several reporter lines and showed that different cell populations identified in scRNA-seq analysis occupied distinct locations in the tissue (Zhu et al., 2022). However, such transgenic reporter analysis is limited in its throughput and thus cannot take full advantage of large-scale molecular information gained by single-cell omics.

Filling this gap is spatial transcriptomics, which spatially maps the expression of hundreds to thousands of genes in a tissue context. Among various types of assays (Moses and Pachter, 2022), hybridization-and imaging-based spatial transcriptomics, such as Multiplexed Error Robust Fluorescence In Situ Hybridization (MERFISH) (Chen et al., 2015) and seqFISH (Eng et al., 2019), can profile gene expression of a tissue section at the single-cell resolution; it is also possible to target bacterial RNAs (Dar et al., 2021). Integration of single-cell multiomics and spatial transcriptomics can provide a spatially-resolved comprehensive molecular cell atlas, but the potential has yet to be explored in the context of host-microbe interactions or plant biology.

In this study, we performed single-nucleus multiome (snMultiome; snRNA-seq + snATAC-seq) and spatial transcriptome (MERFISH) analyses of mature *A. thaliana* leaves infected by both virulent and avirulent bacterial pathogens at four different time points at early infection stages. We used three bacterial pathogens: DC3000, a virulent pathogen that can suppress plant immunity by effectors and toxins; DC3000 AvrRpt2 and DC3000 AvrRpm1, avirulent pathogens that carry effectors recognized by plant NLRs to trigger ETI. Time-resolved single-cell multiomics analysis revealed many previously uncharacterized immune cell populations and potential gene regulatory modules containing TFs, putative CREs, and target genes associated with both disease and resistance. In addition, the integration of snMultiome and MERFISH datasets revealed the spatiotemporal organization of the transcriptome and epigenome with bacterial colonization information. Finally, a web tool that allows users to explore these atlas datasets is available (plantpathogenatlas.salk.edu).

## Results

### Time-resolved snMultiome analysis in plants infected by bacterial pathogens

We generated a time-resolved snMultiome atlas of *A. thaliana* leaves infected by three bacterial pathogens: one is virulent (DC3000), and two are avirulent (AvrRpt2 and AvrRpm1) (Figure 1A). Pathogens were syringe infiltrated at a low dose (OD_600_ = 0.001), where only a subset of plant cells are expected to encounter pathogen cells. We chose four different time points, which are at an early stage of infection, where dynamic transcriptional reprogramming was observed in a previous study with bulk RNA-seq (Mine et al., 2018). We also prepared plants infiltrated with water (mock) and sampled them after 9 h as a control. All samples were collected at the same time of the day (i.e., pathogens were inoculated at different timings) to minimize the influence of circadian rhythms. We developed a protocol to quickly isolate nuclei that minimizes transcriptome and epigenome changes during sample preparation (see Methods). A total of 41,994 cells from 13 conditions passed QC based on both RNA-seq and ATAC-seq data (Figure S1A-D; see Method for QC cutoff). The snATAC-seq analysis detected more ACRs (35,617 ACRs) than a previous bulk ATAC-seq study of immune-activated *A. thaliana* leaves (24,901–27,285 ACRs) (Ding et al., 2021), probably because snATAC-seq could capture cell type-specific signal that could be diluted in bulk ATAC-seq. Peaks associated with a housekeeping gene (*ACTIN2*) were consistently detected across all the samples, supporting the high reproducibility of snATAC-seq data (Figure S1E).

**Figure 1.**
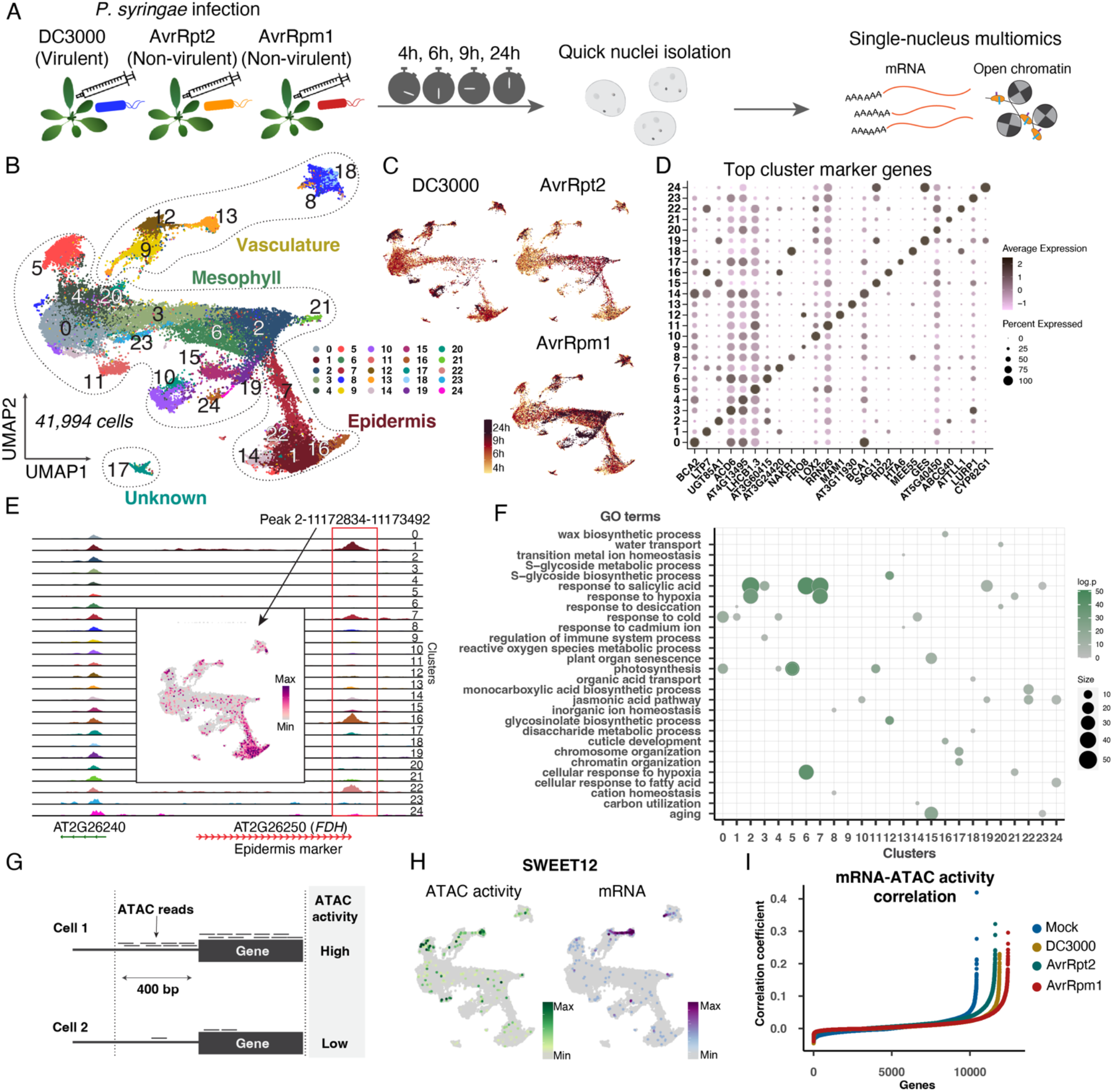
Time-resolved single-nucleus multiomics of *A. thaliana* leaves infected by bacterial pathogens. (A) Schematic diagram of time-course single nucleus multiomics (B) Two-dimensional embedding of nuclei from all samples with uniform manifold approximation and projection (UMAP) based on the transcriptome data. Nuclei are colored by Leiden clusters. (C) UMAP plots for each infection condition. Nuclei are colored by time points. (D) Dot plot showing the top marker genes of individual clusters. A full list is available in Table S1. (E) Cluster-aggregated chromatin accessibility surrounding *FDH*, a known marker gene for leaf epidermis. The overlaid UMAP plot shows the ATAC-seq count on a peak near *FDH* (chromosome 2, position 11172834-11173492) in each nucleus. (F) GO enrichment analysis for marker genes of each cluster. (G) Schematic diagram of the ATAC activity analysis. For each gene, ATAC reads mapped on the gene body or the 400 bp upstream region were aggregated to calculate the score. (H) ATAC activity score (left) and mRNA expression (right) of *SWEET12*, a phloem marker. (I) Pearson correlation coefficients between mRNA expression and ATAC activity for each gene

Dimensionality reduction and clustering were performed independently on snRNA-seq and snATAC-seq data, with snRNA-seq data showing better resolution (Figure 1B; Figures S1F); joint embedding and clustering of snRNA-seq and snATAC-seq showed a similar result with snRNA-seq alone (Figure S1G). Thus, we used the clustering based on snRNA-seq data for the rest of the analysis. Some clusters were enriched with specific infection conditions and time points (Figure 1C; Figure S1H), suggesting that the clustering captured distinct cell states induced by pathogen infection. We identified genes specifically expressed in individual clusters (top markers were shown in Figure 1D), which further clarified the identity of each cluster (major cell type annotations were shown in Figure 1B). Cell types could also be predicted based on ATAC-seq data. For instance, a cluster-specific ACR (peak 2-11172834-11173492) of clusters 1, 7, 14, and 22 was associated with *FIDDLEHEAD (FDH)*, an epidermis marker gene (Figures 1E). Overall, snMultiome data classified cell types/states of *A. thaliana* leaves under pathogen attack. We performed a functional enrichment analysis for each cell cluster (Figure 1F). GO enrichment analysis of marker genes of each cluster revealed that clusters 2 (mesophyll), 6 (mesophyll), and 7 (epidermis) were enriched with defense-related genes, including those involved in the defense hormone salicylic acid (SA) pathway (Peng et al., 2021) (Figure 1F). Notably, these clusters were well-represented in ETI-triggering conditions (AvrRpt2 and AvrRpm1 infection) (Figure 1C; Figure S1H), suggesting that these cells were responding to the immune-triggering pathogens, which is supported by the specific induction of *ICS1*, a key gene for pathogen-induced SA biosynthesis (Wildermuth et al., 2001) (Figure S1I). Furthermore, we observed increased accessibility to chromatin regions upstream of the *ICS1* locus upon infection by the ETI-triggering strains (Figure S1J), which is consistent with a previous bulk ATAC-seq study (Ding et al., 2021). Importantly, defense-active clusters were mostly absent from the mock sample (Figure S1H and S1K), indicating that the infiltration and nuclei isolation processes did not introduce significant artifacts that affect the analysis of plant defense responses at the single-cell level. Together, our results captured heterogeneous and coordinated changes in defense gene expression and chromatin remodeling upon pathogen attack.

To further compare RNA-seq and ATAC-seq data, we calculated the gene activity score by summing ATAC counts on the gene body and 400 bp upstream of the gene, which was used in a previous study (Dorrity et al., 2021) (Figure 1G). Many genes showed correlated mRNA expression and gene activity score as exemplified by *SWEET12* (Figures 1H and 1I), indicating that, in some cases, gene activity score can be used as a proxy for gene expression in *A. thaliana* leaves. However, there were many cases where activity scores did not correlate with gene expression (Figure 1I), demonstrating the need for multimodal analysis.

### Links between gene expression and chromatin accessibility

Our snMultiome data allowed us to directly compare mRNA expression and ACRs to identify potential CREs in different cell types, infection conditions, and time points (Figure 2A). We identified a total of 84,481 significantly correlated (or “linked”) ACR-gene pairs within 500 kb of each gene across cells within each infection condition (Table S2). A majority of links were within a short distance (<400bp; potential promoter regions), but many links were farther away from genes (potential enhancer regions) (Figures 2B-2C). Cluster marker genes (1,774 genes) had more linked ACRs in a moderately distal region (2.5 kb to 5 kb from TSS) compared with non-marker genes (Figure 2D), implying the role of CREs moderately distal from the TSS in cell type/state-specific gene regulation, which is consistent with the observation in a previous study (Tu et al., 2021). We also found that linked ACRs were slightly larger than non-linked ACRs (Figure S2A). Some genes had more than one link (Figure S2B), and we observed that the number of links explains how well the ATAC activity score correlates with gene expression (r^2^=0.64; Figure S2C). Nevertheless, we found that maximum Pearson correlation coefficient values between each gene and linked peaks are a better predictor of correlations between gene expression and gene activity score (r^2^=0.79; Figure S2D).

**Figure 2.**
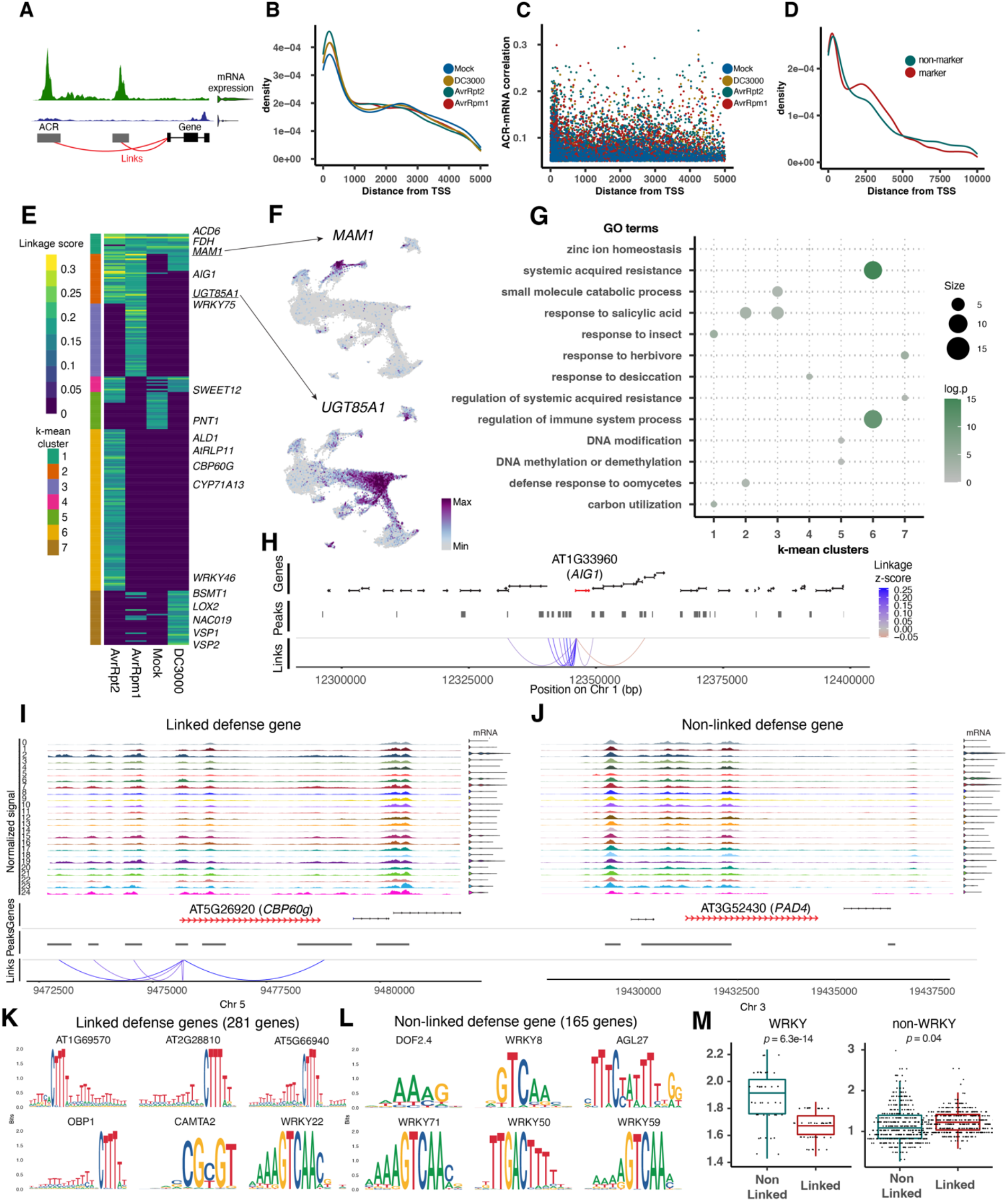
Linking gene expression and chromatin accessibility at the single-cell level. (A) Schematic diagram of linked ACRs and a gene. An accessible chromatin region (ACR) and a gene is “linked” when there is a significant correlation between chromatin accessibility and mRNA expression across individual cells. (B) Density plot showing the frequency of linkages at different distances from the transcription start sites (TSSs). (C) Scatter plot showing the distribution of the linkage score (Pearson correlation coefficient between ACR count and mRNA expression) at different distances from TSSs. (D) Density plot showing the frequency of linkages at different distances from TSS for cluster marker genes and non-marker genes. (E) Heatmap showing the linkage score for genes that showed at least one significant link in at least one of the infection conditions. When a gene had multiple links, the maximum linkage score was shown. The sidebar shows the k-mean cluster annotation. (F) Expression of mRNA encoding *MAM1* and *UGT85A1*. (G) GO enrichment analysis of genes in each k-mean cluster shown in (E) (H) Linkages between ACRs and the pathogen-responsive gene *AIG1*. (I-J) Cluster-aggregated chromatin accessibility surrounding *CBP60g* (I) and *PAD4* (J). Violin plots on the side show aggregated mRNA expression of each gene. (K-L) Top motifs enriched in the promoter regions (2 kb upstream from the TSS) of defense genes (markers of cluster 2, an immune-active mesophyll cell cluster (Figure 1B)) that are linked (K) and not linked (L). (M) Enrichment of WRKY (left) and non-WRKY (right) motifs in the promoter regions of linked or non-linked genes.

We summarized linkage data for each gene by using the maximum Pearson correlation coefficient values (linkage score) (Figure 2E). Genes that showed linkages in all the conditions (Cluster 1 of Figure 2E) were enriched with cell-type marker genes, such as *FDH*, *BCA2*, and *MAM1* (Figures 2E-2F). Genes that showed linkages specifically in ETI-activated conditions (Clusters 2/3/6 of Figure 2E) were enriched with immunity-related genes (Figures 2E and 2G). *AIG1*, which was induced during ETI (Figure S2E), had a large number of links, specifically in the ETI conditions (Figures 2E and 2H). *CBP60g*, a transcriptional regulator of immunity, has multiple ACRs whose accessibility significantly correlates with its mRNA expression (Figure 2I). Genes that showed linkages specifically in DC3000 infection (Cluster 7 of Figure 2E) were enriched with jasmonic acid (JA)-related genes (“response to herbivore”; Figure 2G), consistent with the ability of DC3000 to activate plants’ JA pathway using the toxin coronatin and effectors to suppress plant immunity (Nobori et al., 2018a). These results indicate that coordinated reprogramming of chromatin accessibility and gene expression is a key feature of leaf development, immunity, and exploitation by virulent pathogens.

### Gene regulation independent from chromatin remodeling

Although we identified many ACRs tightly associated with defense genes, we wondered how prevalent such links are. Interestingly, we found that 37% (165 out of 446; Table S4) of defense genes (marker genes of immune-active mesophyll clusters 2 and 6) did not show any significant link with ACRs. Such non-linked defense genes often harbor constitutively opened chromatin (Figure 2J), despite dramatic changes in gene expression (Figure 2J, violin plot), suggesting that there is an additional layer of gene regulation, such as the expression of upstream TFs. Thus, we performed motif enrichment analysis for ACRs flanking non-linked defense genes (within 2 kb from TSS). The analysis identified significant enrichment of WRKY motifs (Figures 2K and 2L). Interestingly, WRKY motifs were more enriched in ACRs near non-linked defense genes than linked defense genes (Figure 2M). This pattern was not observed for other TF motifs (Figure 2M). Thus, expression and/or binding of WRKY TFs may provide an additional layer of regulation to defense genes with constitutively open chromatin.

### Identification of TF-ACR-gene modules associated with pathogen infection in single cells

We sought to identify TFs that regulate candidate CREs. We performed motif enrichment analysis for “linked” ACRs specific to an immune-active mesophyll cluster (cluster 2) compared with a non-immune-active mesophyll cluster (cluster 0) and found the enrichment of many TFs known to be involved in immunity, including WRKYs, CAMTA2, and ANAC050 (Figure 3A). The same analysis comparing the immune-active epidermis (cluster 7) and non-immune-active epidermis (cluster 1) showed the enrichment of a similar set of TF motifs, whereas comparison between the non-immune-active epidermis and mesophyll showed the enrichment of different TF motifs (Figures 3B). Notably, in this cell type comparison, *EDT1* (also known as *HDG11*), a homeodomain (HD)-START TF, whose binding motif was enriched in the epidermis, has previously been shown to function in the patterning of the epidermis and drought tolerance (Yu et al., 2008), indicating that the analysis can capture biologically relevant TFs in target cell populations. We extended this analysis to marker ACRs for all the clusters and identified cluster-specific motif enrichment (Figure 3C). These results indicate that different cell types and states employ both shared and unique gene regulation via TF-DNA binding and will accelerate the identification of TFs with cell type/state-specific function.

**Figure 3.**
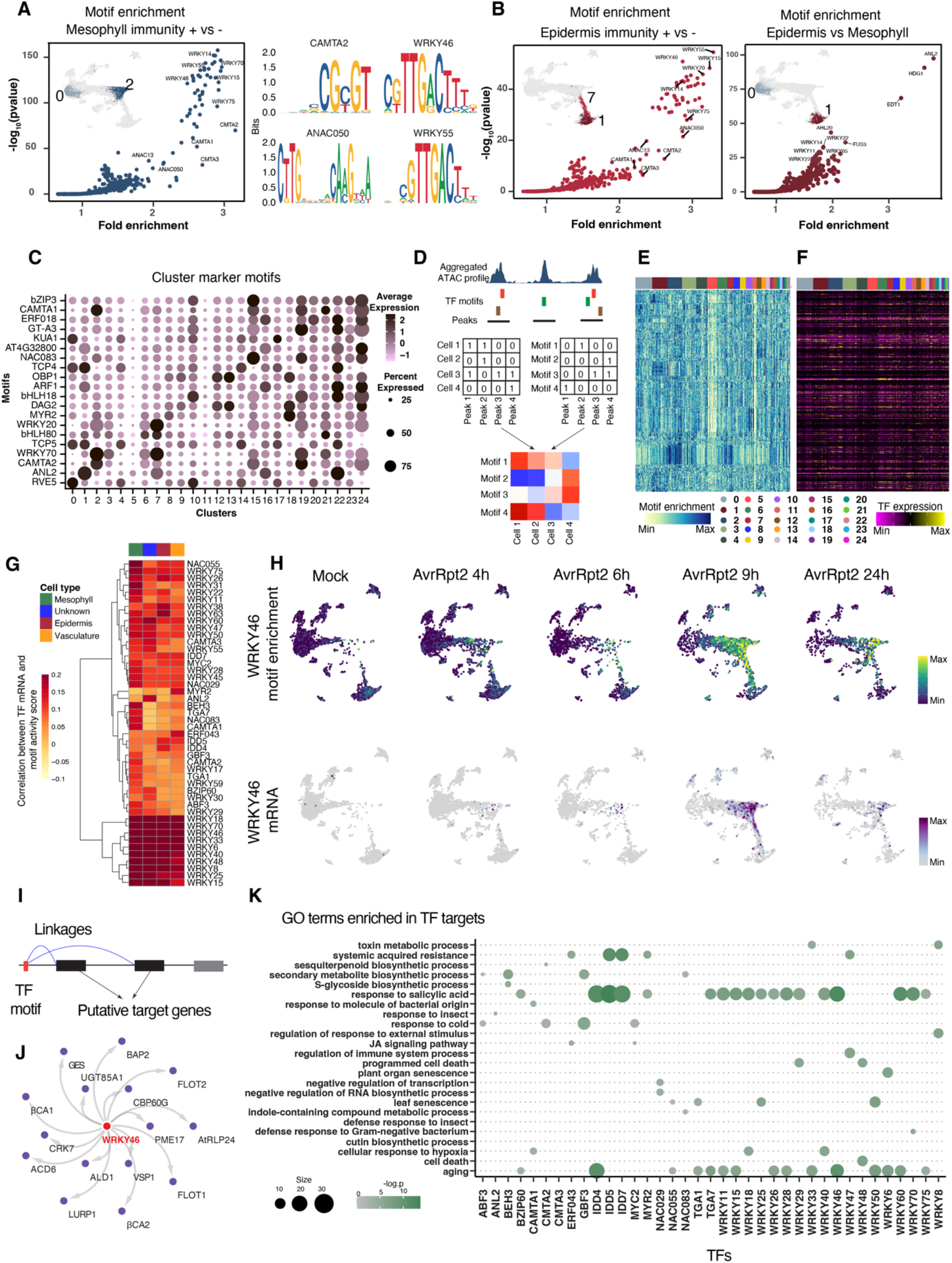
Identification of TF-ACR-gene modules. (A) (Left) Scatter plot showing motif enrichment in cluster 2 (immune-active mesophyll) compared with cluster 0 (non-immune-active mesophyll). (Right) Selected motifs enriched in cluster 2. (B) Scatter plot showing motif enrichment in (left) cluster 7 (immune-active epidermis) compared with cluster 1 (non-immune-active epidermis) and (right) cluster 1 compared with cluster 0. (C) Top motifs enriched in each cluster. A full list is available in Table S5. (D) Schematic diagram of motif enrichment score analysis. ChromVAR (Schep et al. 2017) was used to generate the “cell x motif matrix” by combining the “cell x peak (ACR) matrix” and the “peak x motif matrix.” (E-F) Heatmaps showing (E) enrichment scores of 465 motifs and (F) expression of corresponding transcription factors (TF) in each nucleus. The top bars show cluster annotation defined in Figure 1B. (G) Heatmap showing Pearson correlation coefficient between motif enrichment scores and mRNA expression of the corresponding TFs in each cell type (shown in the top bar). (H) Motif enrichment scores (top) and mRNA expression (bottom) of *WRKY46*. (I) Schematic diagram of target gene analysis of TF motifs. Genes that are significantly linked to any ACR containing a motif of interest were defined as putative targets. (J) Predicted target genes of *WRKY46*. (K) GO enrichment analysis of genes predicted to be targeted by TFs shown in (G).

To analyze motif accessibility at the single-cell level–– not the cluster level––we calculated the motif enrichment score for each cell using ChromVAR (Schep et al., 2017) (Figure 3D), which revealed heterogeneous accessibility to 465 motifs between and within clusters (Figure 3E). We then asked whether the TF motif enrichment score correlates with TF expression, which also showed heterogeneous expression (Figure 3F). We found a number of TFs that show a high correlation between motif enrichment scores and their mRNA expression in different cell types (Figure 3G). For instance, WRKY46 mRNA expression and accessibility to WRKY46 binding sites overlapped well mainly in immune active mesophyll and epidermal cells (Figure 3H), suggesting that the WRKY46 regulon plays a key role during immune responses in these cell types. By searching genes that are linked with ACRs containing WRKY46 binding sites (Figure 3I), we identified potential target genes of WRKY46, including many known defense-related genes (Figure 3J), consistent with the known function of WRKY46 in the SA pathway and defense against *P. syringae* (Hu et al., 2012). Finally, we performed target gene prediction for all the TFs shown in Figure 3G and found that many TFs target genes related to plant defense (Figures 3K and S3A). For instance, our analysis predicted that bZIP60, TGA1, and TGA7 commonly regulate *WRKY18* and other defense-related genes (Figures 3K and S3A). *IDD4/5/7* were also predicted to regulate various defense genes, including *CBP60g* (Figures 3K and S3A). Expression of *IDD4* was decreased upon the infection of AvrRpt2 (Figures S3B), which is consistent with a previous study that showed that IDD4 negatively regulates plant immunity (Völz et al., 2019). In summary, we identified a wealth of TF-ACR-gene modules that characterize different cell types and pathogen-responding cells.

### Fine dissection of immune-activated cell populations

Although the major clusters captured immune active cells in mesophyll and epidermis (Figure 1F), clusters for other cell types, such as vasculature, contained both immune-active and non-active cells, likely due to strong developmental signatures. To identify cell type-specific immune responses, we performed a second round of clustering for each major cluster, which resulted in 231 sub-clusters with diverse transcriptome patterns (Figure 4A). Sub-clustering of major cluster 8, which showed strong expression of Phloem Companion Cells (PCCs) markers (Figure S4A), resulted in 11 sub-clusters, some of which were enriched with immune genes (Figure 4B).

**Figure 4.**
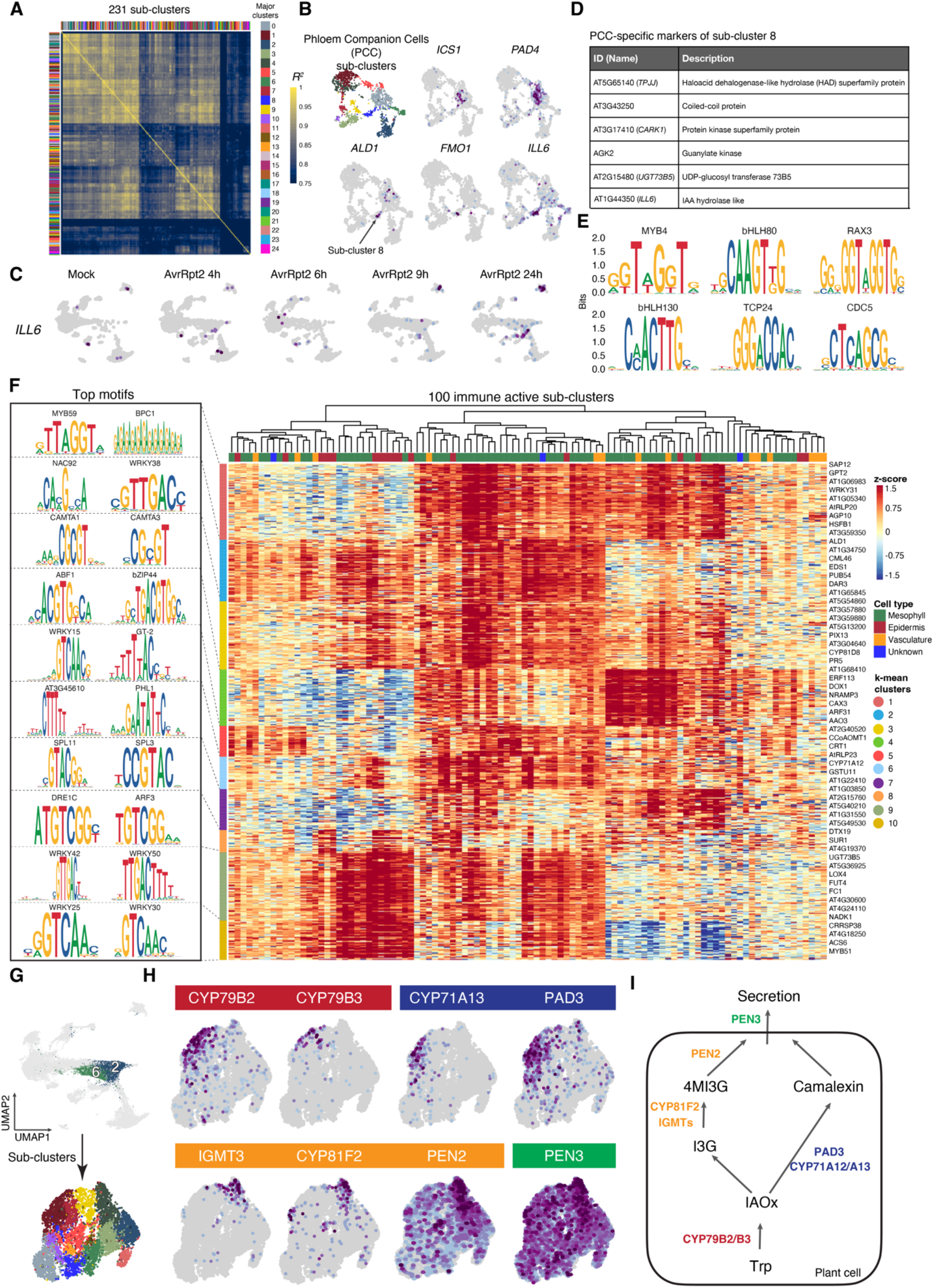
Identification of cell type-and cell state-specific immune responses. (A) Heatmap showing pair-wise correlation of pseudobulk transcriptomes between sub-clusters of individual clusters. The top and side bars show major cluster labels. (B) Sub-clustering of cluster 8 (phloem companion cells). Defense-related genes showing sub-cluster-specific expression were shown. (C) Expression of *ILL6* upon infection of AvrRpt2 in time course, showing specific induction in cluster 8. (D) A list of genes specifically expressed in sub-cluster 8-8. (E) Top motifs enriched in the marker genes of sub-cluster 8-8. (F) Heatmap showing normalized expression of genes associated with immunity (rows; see Method for gene selection) across top 100 sub-clusters that express these genes (columns; see Method for cluster selection). Names of representative genes are shown on the right. The top bar shows the cell type of origin. The sidebar shows k-mean cluster annotation. Top two motifs enriched in the accessible promoter regions (2 kb upstream from the TSS) of genes in each k-mean cluster were shown. (G) Schematic diagram of sub-clustering of clusters 2 and 6 (immune-active mesophylls). (H) Expression of genes involved in different steps of the biosynthesis and secretion of Tryptophan (Trp)-derived secondary metabolites shown in (I). (I) Simplified schematic diagram of the biosynthesis and secretion of Trp-derived secondary metabolites.

Key genes for systemic acquired resistance (SAR), *ALD1* and *FMO1*, were specifically expressed in PCC sub-cluster 8 (Figure 4B), implying that a subset of PCC cells contributes to sending the long-distance signal to systemic leaves. Notably, we did not observe strong expression of *ALD1* and *FMO1* in other vasculature clusters (Figure S4B). We found *ILL6* specifically induced in PCC sub-cluster 8 (Figures 4B-4C); *ILL6* was previously shown to be involved in SAR (Khan et al., 2021), implying that this cell population plays a role in SAR, and *ILL6* may be a PCC specific SAR regulator. We then systematically searched for genes specifically enriched in PCC sub-cluster 8 but not in other major clusters and identified many such genes (Figure 4D). These genes may also be involved in the specific function of PCC cells during SAR. Among them, *UGT73B5* has been shown to contribute to ETI (Langlois-Meurinne et al., 2005). The accessible promoter regions of the unique marker genes of PCC sub-cluster 8 were enriched with binding sites for MYB4, bHLH80, and RAX3 (MYB84) (Figure 4E), which may play important roles in SAR.

We screened all 231 sub-clusters and selected 100 cell populations with immunity signals based on the expression of known immune genes and GO enrichment analysis (see Methods for detail). These putative “immune cell populations” showed varying gene expression and motif enrichment patterns, suggesting that there are multiple immune states in individual developmental cell types (Figure 4F). For example, some genes enriched with WRKY motifs (clusters 9 and 10 of Figure 4F) were particularly highly expressed in sub-cell populations of the mesophyll and epidermis, whereas other genes enriched with ABF1 and bZIP44 motifs (cluster 4 of Figure 4F) were highly expressed in other sub-cell populations of the mesophyll and vasculature. Mesophyll and epidermal cells tend to show stronger immune responses compared with vasculature cell types and cells with unknown identities (Figure 4F and Figure S4E). Interestingly, cell populations from different cell types can take a similar immune cell state (Figure 4F), suggesting that some immune gene regulatory mechanisms are shared among multiple developmental cell types.

As an example of cell population-specific immune responses, we analyzed Tryptophan-derived defense-related secondary metabolite pathways in immune active mesophyll cells (clusters 2 and 6) (Figure 4G). We found a distinct expression of genes involved in different steps/pathways of the biosynthesis/secretion of the defense metabolites camalexin and indole glucosinolate (Figures 4H-4I), suggesting there is compartmentalization in the activation of defense pathways that can potentially compete resources (Tryptophan).

### Time-resolved spatial transcriptomics reveals immune sub-populations

To validate cell populations identified in snMultiome analysis and analyze gene regulatory modules in the actual tissue context, we performed MERFISH on tissue sections of infected leaves (Figure 5A). MERFISH is a massively multiplexed single-molecule imaging that can detect hundreds or thousands of RNA species at the single-cell resolution (Chen et al., 2015) (Figure S5A). We curated 500 target genes (Table S6), including (1) leaf cell type markers, (2) genes involved in various processes such as immunity, hormone pathways, and epigenetic regulation, and (3) a variety of TFs. In addition to MERFISH, we performed standard smFISH on the same tissues targeting *ICS1* (see Methods); we also targeted bacterial genes to locate bacterial cells in the tissue section (see Methods). We profiled leaves infected by AvrRpt2 at each of the four time points, matching the snMultiome experiments, as well as the mock condition (Figure 5A). The spatial localization of 500 genes was decoded after the combinatorial smFISH imaging, detecting millions of transcripts per sample (Figure 5B).

**Figure 5.**
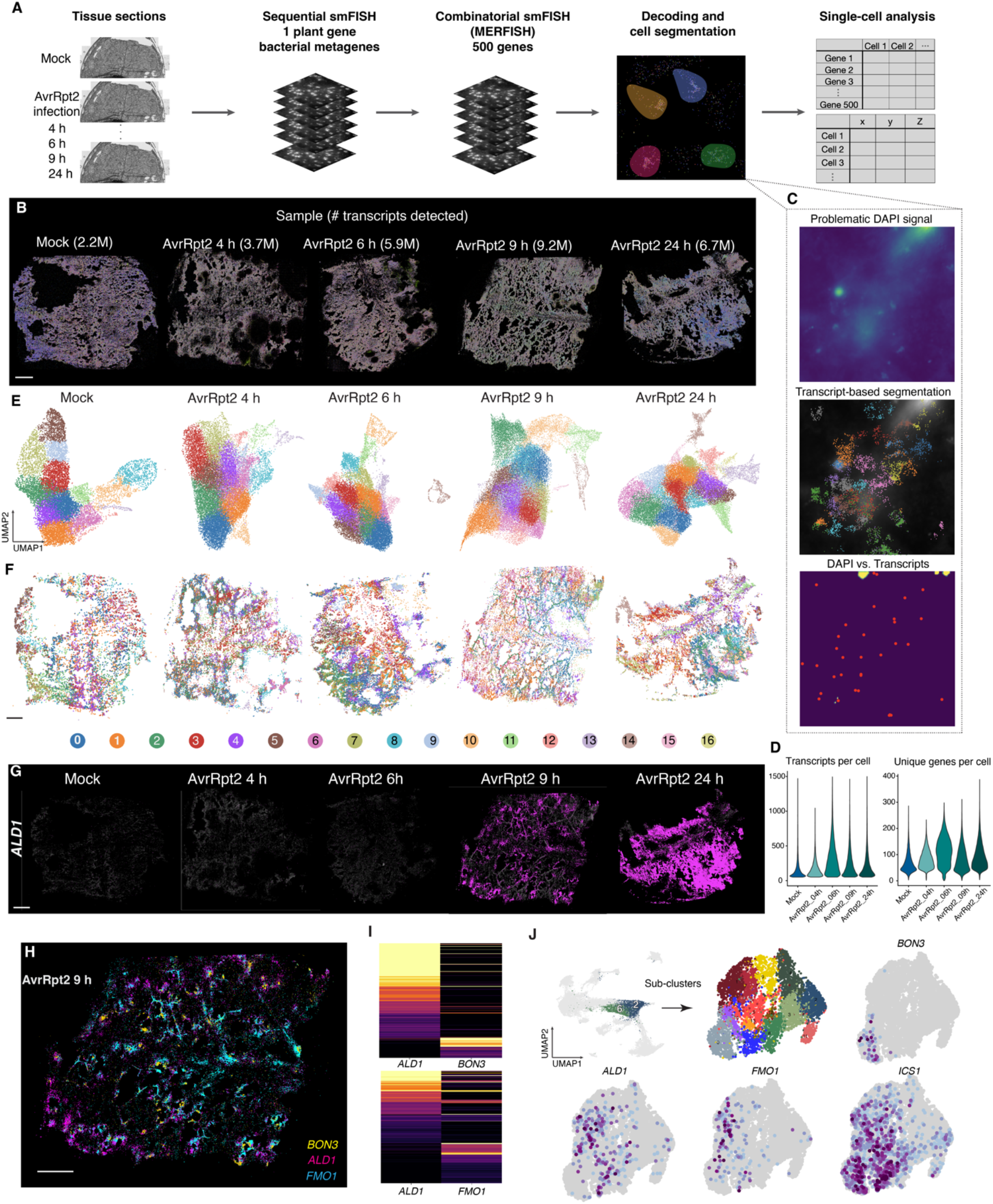
Time-resolved spatial transcriptomics in pathogen-infected leaves. (A) Schematic diagram showing MERFISH experiments. (B) Two-dimensional plots of all the transcripts detected by MERFISH in each sample. The number of transcripts is shown. (C) (Top) A field of view (FOV) that shows obscure DAPI nuclei staining signal. (Middle) Transcript-based segmentation in the same FOV (see Method). Transcripts were colored by assigned cells. (Bottom) Centroids of cells detected by the transcript-based segmentation (red dots) and the result of failed DAPI-staining-based segmentation (yellow region). (D) Violin plots showing the number of transcripts per cell (left) and unique genes per cell (right) detected in each MERFISH sample. (E) UMAP embeddings of cells in each sample based on the expression of 500 genes detected by MERFISH. Cells are colored by *de novo* Leiden clusters. (F) Spatial mapping of Leiden clusters in each sample, using the same color scheme as (E). (G) Spatial expression pattern of *ALD1* detected by MERFISH in each sample. (H) Spatial expression pattern of *ALD1*, *FMO1*, and *BON3* at 9 h post-infection (hpi). (I) Heatmaps showing expression of *ALD1*, *FMO1*, and *BON3* in each cell at 9 hpi. (J) Expression of *ALD1*, *FMO1*, *BON3*, and *ICS1* in the sub-clusters of clusters 2 and 6 in the snRNA-seq data. (B, F, G, and H) Scale bars = 200 µm.

A key step for the single-cell analysis of MERFISH data is cell segmentation (Figure 5A). We found that the standard segmentation approach with nuclei (DAPI) and cytoplasmic (poly(A)) staining did not provide high-quality segmentation results (Figure 5C). We, therefore, implemented a new segmentation approach based on the transcript distribution, which successfully segmented cells (see Methods; Figure 5C; Figures S5B-D). After segmentation, transcripts were assigned to cells, resulting in a cell-by-gene matrix with each cell having its spatial coordinate (Figure 5A). Overall, we detected a median of 161 transcripts and 79 genes from a total of 121,998 cells from five leaf sections (mock and four time points after AvrRpt2 infection) (Figure 5D). We performed *de novo* cell clustering in each sample, consistently identifying 15 to 17 clusters (Figure 5E). These clusters were spatially mapped, revealing spatially organized cell populations (Figure 5F). For instance, a *de novo* MERFISH cluster captured vasculature cells in a tissue section (Figure S5E). Moreover, MERFISH captured induction of the defense gene *ALD1*, with temporal dynamics consistent with our snMultiome data (Figure 5G; Figure S5F). Thus, snMultiome and MERFISH validate each other’s results, and MERFISH analysis faithfully captured spatial gene expression on pathogen-infected leaf tissue in a time course.

MERFISH allowed us to investigate the spatial expression of many pathogen-responsive genes. We found that *BON3* was induced in highly restricted areas in the tissue upon infection by AvrRpt2 (at 9 h post-infection or hpi) (Figure 5H). The expression pattern was distinct from other immune genes, such as *ALD1* and *FMO1* (Figure 5H). *BON3* and *ALD1* were expressed in different but adjacent cells (Figures 5H-5I), implying distinct functions of these cells and potential cell-cell communication between these cells. *FMO1*, which functions immediately downstream of ALD1 for the production of NHP (a critical molecule for SAR), tended to be co-expressed with *ALD1* more than *BON3*, but there are cells that express only one of these genes (Figures 5H-5I), raising the possibility that multiple cells may be involved in completing the SAR pathway. Our sub-clustering analysis of snMultiome data confirmed the pattern observed in MERFISH (Figure 5J). Interestingly, *ICS1* was detected in cells that express either of the three genes (Figure 5J), suggesting that SA biosynthesis plays a role in broader cell populations.

### Integration of snMultiome and spatial transcriptome data

We found that cell populations defined by MERFISH and snRNA-seq were in good agreement (see Method; Figure 6A; Figure S6). The correspondence between two unimodal datasets motivated us to integrate both data types aiming to spatially map the entire snMultiome data. We integrated snMultiome (five conditions matching to the MERFISH samples; Figure 6B) and MERFISH data using the shared 500 genes, and cluster labels defined by snMultiome were transferred to the MERFISH cells (Figure 6B; see Method). The integrated UMAP showed a good representation of cells derived from the two assays (Figure 6C). Based on the data integration, we spatially mapped clusters defined in the snMultiome data. Major cell types defined by snMultiome were successfully mapped on the expected regions in a MERFISH tissue (Figure 6D), indicating a successful data integration. Note that although the section was in the middle of the leaf, some epidermal cells were included since the section was not completely flat. We found that the snMultiome cluster 16 (or cluster 17 in Figure 1B), which has an unknown identity enriched with histone proteins, is sparsely distributed across the tissue (Figure 6E), which is confirmed by the spatial expression patterns of the top marker genes *HTA6* and *MET1* (Figure 6F). The identity and function of this cell population still remain unknown, but S phase marker genes, *HIS4* and *TSO2* (Menges et al., 2002), were enriched in this cluster (Figure S6C and S6D), implying that cells in this cluster may undergo DNA synthesis. Notably, sub-populations of this cell population showed immune responses upon pathogen attack (Figure 4F), providing a potential opportunity to study cell cycle-specific immune responses. The distribution of an immune active cluster (cluster 6) was consistent with the spatial expression patterns of *ALD1* (Figures 6E and 5G). Overall, the successful integration of MERFISH and snMultiome allowed us to explore the spatial distribution of cell populations defined in snMultiome analysis.

**Figure 6.**
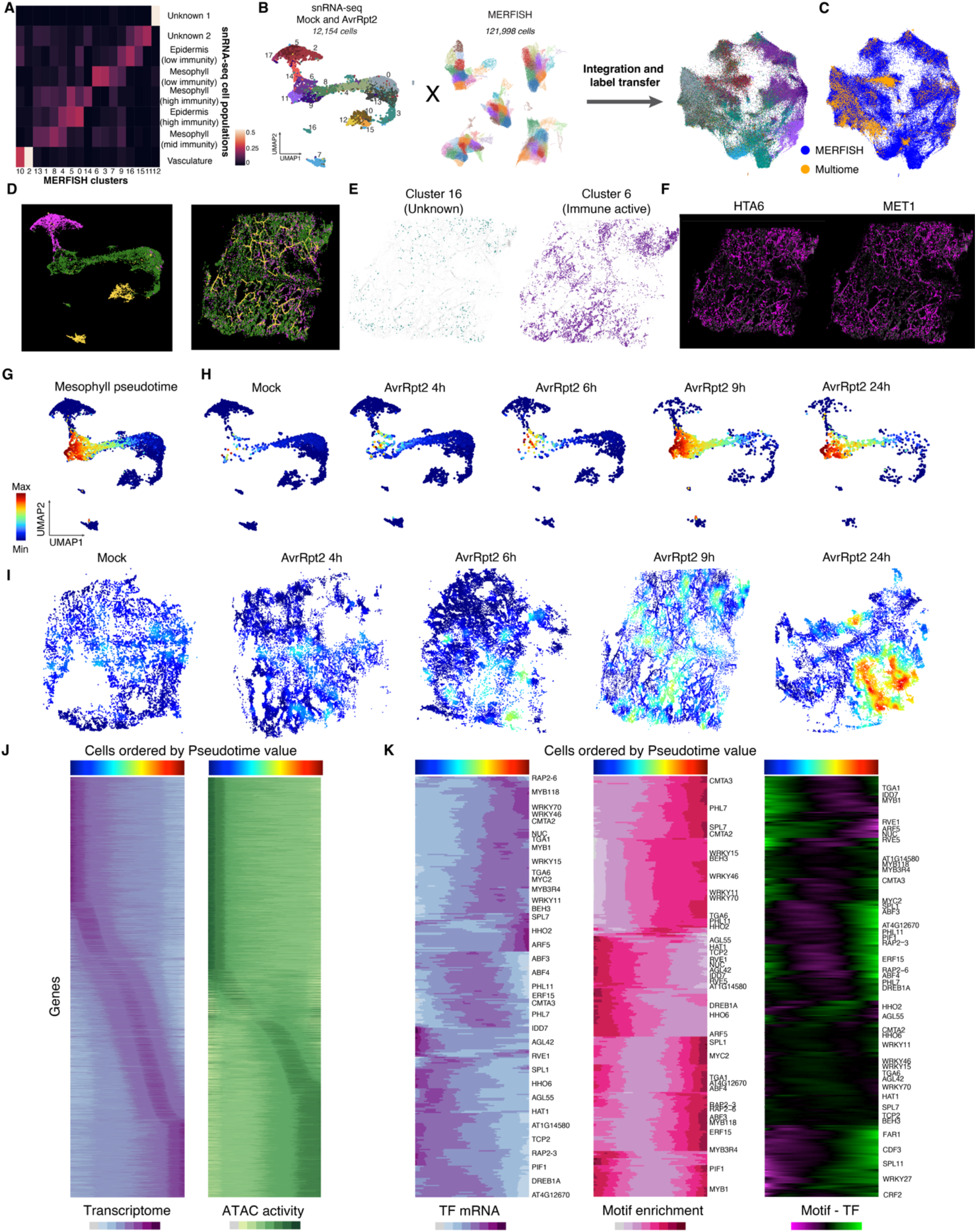
Integration of snMultiome and spatial transcriptome data. (A) Proportion of *de novo* MERFISH clusters (9 h post-infection (hpi), shown in Figure 5E) assigned to cell populations defined by snMultiome (RNA data) based on data integration (see Method). (B) Integration of snMultiome data (mock and AvrRpt2-infected; five samples) and the MERFISH data shown in a UMAP. Nuclei/cells are colored by cluster labels, which were transferred from snRNA-seq to MERFISH (see Method). (C) The integrated UMAP colored by assay types. (D) (Left) The same UMAP as (B) colored by major cell types: epidermis (magenta), mesophyll (green), and vasculature (yellow). (Right) Spatial mapping of snMultiome cells colored by major cell types. AvrRpt2 9 hpi sample was used. (E) Spatial mapping of clusters defined by snMultiome based on data integration and label transfer. Clusters of unknown identities (cluster 16) and immune-active cells (cluster 6) were selected. (F) Spatial expression of *HTA6* and *MET1*, marker genes for cluster 16. (G) Pseudotime values calculated for mesophyll cells in the snRNA-seq data. (H) UMAP plots showing pseudotime values in cells from each time point. (I) Spatial mapping of pseudotime values based on data integration and label transfer. (J) Heatmaps showing transcriptome and ATAC activity (Figure 1G) of corresponding genes for aggregated cells along the pseudotime trajectory. (K) Heatmaps showing TF mRNA expression (Figure 3E), motif enrichment scores of corresponding TFs (Figure 3F), and the difference between them for aggregated cells along the pseudotime trajectory. For each heatmap, the same set of genes is labeled.

### Mapping spatiotemporal dynamics of plant immune responses

To understand the temporal dynamics of immune responses, we applied pseudotime analysis to our snMultiome data. Pseudotime analysis aligns cells as a trajectory based on their gene expression, which is commonly used for inferring developmental trajectories (Saelens et al., 2019). We hypothesized that by applying pseudotime analysis to a single developmental cell type with various immune states, we could better model the temporal dynamics of the heterogeneous infection/immune responses. We calculated pseudotime scores for mesophyll cells (Figure 6G; see Methods). Remarkably, the distribution of predicted pseudotime scores in each sample was consistent with what is expected from the real sampling time point (Figure 6H), i.e., cells with low pseudotime scores were enriched in early time points, and cells with high pseudotime scores emerged at later time points, indicating the successful modeling of the temporal dynamics of immune responses. Cells with a wide range of pseudotime scores coexisted at 9 and 24 hpi, suggesting that cellular immune responses are asynchronous in infected leaves. Are the asynchronous immune states spatially coordinated? To understand the spatial distribution of heterogeneous immune states, we spatially mapped the pseudotime score using label transfer from snMultiome to MERFISH data (Figure 6I). The spatial mapping of the pseudotime score revealed immune active areas distributed in pathogen-infected leaf tissues in a restricted manner (Figure 6I). These immune-active areas expanded over time and appeared to merge at 24 hpi (Figure 6I). Notably, immune states also dynamically change over time within each immune active area, with “older” immune-active cells (higher pseudotime score) being surrounded by “younger” immune-active cells (lower pseudotime score) (Figure 6I). The results imply that the “oldest” immune-active cells are likely plant cells that had direct contact with pathogen cells at early time points, and immune responses spread to surrounding cells via cell-cell communications over time.

The analysis of transcriptome and ATAC activity (Figure 1J) for each gene along the pseudotime immune trajectory revealed continuous and dynamic changes within the transcriptome and epigenome (Figure 6J). To gain insights into gene regulatory mechanisms along pseudotime, we focused on TF expression and motif enrichment (Figure 6K). We found some WRKY and MYB TFs induced at the early stages of immune activation, whereas other TFs, such as SPL7, HHO2, and ARF5, were induced later. The motif enrichment scores of corresponding WRKYs showed temporal dynamics similar to mRNA expression of the TFs, suggesting concordant TF expression and epigenomic changes of WRKY modules (Figure 6K). Other TFs showed earlier or later changes of motif accessibility compared with mRNA expression (Figure 6K), suggesting varying temporal dynamics in the gene regulatory landscape.

### Spatial mapping of transcriptome and epigenome

Integration of MERFISH and snMultiome data allowed us to spatially impute the whole transcriptome and chromatin accessibility information. Imputed *ALD1* expression (based on snMultiome) accurately predicted real spatial expression of *ALD1* (based on MERFISH) (Figure 7A), indicating the data imputation is highly accurate. Moreover, imputed *ICS1* expression accurately predicted real spatial expression of *ICS1* (based on smFISH) (Figure 7B). We also spatially imputed ATAC activity scores of *ICS1* and *ALD1*, which showed consistent patterns with mRNA expression (Figure 7C). The motif activity of HsfB2b (HEAT SHOCK TRANSCRIPTION FACTOR B2b) was predicted to be high in the immune active regions of a leaf at 24 hpi (Figure 7D), which was consistent with the mRNA expression pattern of HsfB2b validated by MERFISH (Figure 7D). The same trend was observed for WRKY31 (Figure S7A). Overall, these results indicate that the spatial data imputation of transcriptome and epigenome was accurate.

**Figure 7.**
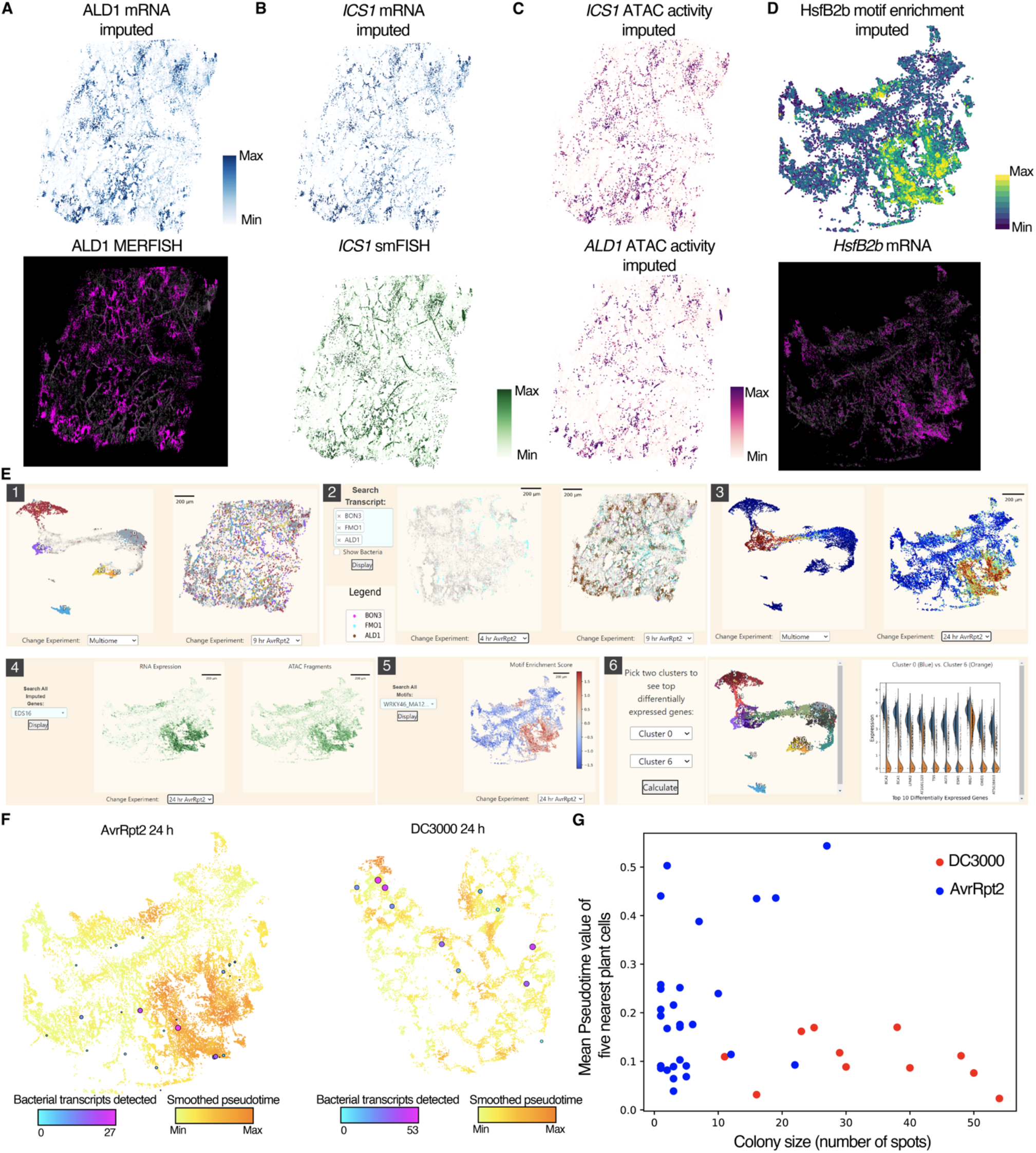
Spatial mapping of whole transcriptome and epigenome. (A) (Top) Imputed mRNA expression of *ALD1*. (Bottom) mRNA expression of *ALD1* with MERFISH. (B) (Top) Imputed mRNA expression of *ICS1*. (Bottom) mRNA expression of *ICS1* with smFISH. (C) Imputed ATAC activity (Figure 1G) of *ICS1* (top) and *ALD1* (bottom). (D) (Top) Imputed motif enrichment scores of HsfB2b. (Bottom) mRNA expression of *HsfB2b* by MERFISH. (E) Overview of a database that hosts the atlas dataset. (1) Spatial mapping of clusters defined by snMultiome. (2) Spatial mapping of 500 genes targeted by MERFISH (3) Pseudotime analysis. (4) Spatial imputation of mRNA expression and ATAC activity. (5) Spatial imputation of motif enrichment. (6) Differential gene expression analysis between two clusters. (F) Spatial mapping of bacterial transcripts detected with smFISH in plants infected by AvrRpt2 (left) and DC3000 (right) at 24 h post-infection (hpi). Pseudotime values (Figure 6H) are also visualized in the background. (G) Scatter plot showing the number of bacterial transcripts (x-axis) and averaged pseudotime values of five nearest neighbor plant cells (y-axis) for each bacterial colony in plants infected by AvrRpt2 (blue) and DC3000 (red) at 24 hpi.

To facilitate the exploration of our data, we imputed all 252,99 transcripts detected in snMultiome and 465 motif enrichment scores on the five tissues used in MERFISH experiments and made the data available on a data browser (plantpathogenatlas.salk.edu). The data browser also hosts all the MERFISH transcripts and spatial mapping of snMultiome clusters and pseudotime scores (Figure 7E).

### Spatial co-mapping of plant responses and pathogen distribution

Finally, we sought to investigate whether spatially restricted immune gene expression can be explained by the distribution of bacterial cells. smFISH targeting bacterial metagenes (19 highly expressed genes) detected bacterial colonies at 24 hpi, which was overlaid with the spatial map of the pseudotime score (Figure 7F). As a control, we performed another MERFISH experiment using an *A. thaliana* leaf infected by the immune-suppressive pathogen DC3000 at 24 hpi and performed the same analysis (Figure 7F and Figure S7B). We observed the distribution of the immune-triggering strain AvrRpt2 overlapped with tissue regions with high pseudotime scores (immune heightened) in contrast to the immune-suppressive DC3000 (Figure 7E- 7F). We confirmed this observation by quantitatively analyzing the neighboring plant cells of individual bacterial colonies (Figure 7G). Taken together, the results suggest that immune active regions defined by the pseudotime analysis indeed interacted with the ETI-triggering pathogen; we also captured potential immune suppression by the virulent DC3000. Interestingly, some AvrRpt2 colonies were not associated with an immune active region, implying that successful ETI activation might require more than the presence of ETI-triggering pathogens.

## Discussion

Plants do not have specialized immune cells, but virtually every cell can mount immune responses. However, the diversity and function of cellular immune states are not fully understood. Our time series snMultiome and spatial transcriptome atlas of pathogen-infected *A. thaliana* leaves revealed various plant cell populations with distinct gene regulatory principles and the spatiotemporal gradient of transcriptome and epigenome changes during immune activation and disease progression. Our snMultiome and spatial transcriptome data using the same conditions matched well, suggesting the high reproducibility of our datasets.

We tested ETI responses triggered by two different effectors, AvrRpt2 and AvrRpm1, and observed similar responses between them, with AvrRpm1 triggering faster responses, which is consistent with a previousstudy (Mine et al., 2018). The two effectors are both recognized by CC-type NLRs, RPS2 and RPM1, respectively (Mindrinos et al., 1994; Bent et al., 1994; Grant et al., 1995). Immune activation by effector-bound RPS2 and RPM1 is considered to be dependent on the oligomerization of itself (RPM1) or helper NLRs (in the case of RPS2), which forms membrane-localized cation channels that let Ca^2+^ into the cell to trigger immune responses (Jacob et al., 2021). Similar single-cell transcriptome patterns between the two ETI responses tested in this study might reflect a common activation mechanism of ETI. It will be interesting to study how other types of NLRs, such as Toll/interleukin-1 receptor (TIR) domain-containing NLRs (Lapin et al., 2022), activate ETI at the single-cell level.

We also identified single-cell plant responses specific to bacterial strains. Some clusters were enriched in plants infected by AvrRpm1 at 24 hpi (clusters 23 and 24) (Figure S1H). Since AvrRpm1 ETI is faster than AvrRpt2 ETI, similar cell states may emerge in AvrRpt2 infected tissues at later time points. Cluster 24 marker genes were enriched with JA pathway (Figure 1G), which is known to be activated upon wounding. It is possible that these cells are related to cell death caused by ETI activation. Clusters 10 (mesophyll) and 22 (epidermis) also highly expressed genes related to JA pathway (Figure 1G) and enriched with cells derived from plants infected by the virulent pathogen (Figure S1H). This may reflect plant responses to pathogen-derived molecules that activate the JA pathway, such as coronatine and effectors (Nobori et al., 2018a). Detailed analysis of these clusters would provide insights into ETI and pathogen virulence.

We identified various immune states based on gene expression (Figure 4F). Interestingly, in most immune states, we found cell populations from multiple developmental cell types (Figure 4F). Also, TFs that showed a high correlation with motif accessibility were shared among developmental cell types (Figure 3G). These observations imply that different developmental cell types can confer shared immune responses with common gene regulatory mechanisms, although the roles of these cell populations may be different. We also found cell type-specific immune responses as exemplified by the identification of PCCs with increased expression of SAR genes (Figure 3B). As SAR requires a mobile signal that travels from locally infected leaves to systemic leaves via vasculature, it is possible that these PCCs play a unique role in SAR.

Our snMultiome data allowed us to investigate the relationships between individual ACRs and genes, revealing many putative TF-ACR-gene modules, including known immune pathways. The results can be used for discovering previously uncharacterized immune genes (Figure S3A); novel defense-related CREs can also be identified by *de novo* motif analysis. Temporal dynamics of TF expression and accessibility to known TF motifs varied (Figure 6J), implying there are different modes of gene regulation. For instance, some TFs may open chromatin by themselves (pioneer TFs), whereas other TFs may be induced after chromatin states are primed by other factors. Further research is needed to understand the mechanism by which chromatin accessibility changes upon pathogen infection.

We found some immune genes were “not linked” and associated with constitutively opened chromatin, while their expression was tightly regulated upon pathogen attack (Figure 2J). The upstream regions of such non-linked immune genes were enriched with WRKY binding motifs (Figure 2L and 2M), implying that tight control of WRKY TFs, which can act as transcriptional activators or repressors, may regulate downstream genes. Constitutively accessible chromatin may facilitate the quick binding of TFs to the DNA for rapid induction of defense genes. It is also possible that non-linked genes are regulated by other epigenetic modifications. It has been shown that DNA methylation could inhibit the binding of WRKY TFs to DNA by steric hindrance (Charvin et al., 2022; O’Malley et al., 2016). It would be interesting to profile DNA methylation at the single-cell level to see if there is such gene regulation. Recently developed methods enable simultaneous single-cell profiling of transcriptome, chromatin accessibility, and DNA methylation from the same cells (Luo et al., 2022). Such “single-cell triple-omics” technologies will further advance our understanding of gene regulation in plants.

MERFISH revealed that *BON3* was induced in highly restricted areas in ETI-triggering tissue (Figure 5H). This information combined with the pseudotime prediction (Figure 6G and 6H) and *BON3* expression patterns (Figure S6B) in snRNA-seq data implies that *BON3*- expressing cells are early responders to pathogen invasion, highlighting the power of combining single-cell and spatialomics analyses. Spatial mapping of pseudotime information modeled how immune responses might spread from cells that recognized pathogen molecules and revealed that cells in different immune states co-exist at a single time point in a spatially organized way (Figure 6I). The model created by the pseudotime analysis was well supported by our time course data, but it would be important to analyze further the temporal dynamics of immune responses with more time points or live imaging. Recent studies employed live cell imaging of transcription dynamics based on RNA labeling (Alamos et al., 2021; Hani et al., 2021). A similar approach could be used to study immune-activated cells.

Finally, this study provides a resource to explore various previously uncharacterized cell populations associated with disease and resistance with spatial and temporal information and potential regulatory mechanisms. Our database can be used as a tool for hypothesis generation and hypothesis testing and will catalyze new discoveries of molecular mechanisms underlying plant-microbe interactions at unprecedented resolution.

### Limitations of the study

Simultaneous spatial mapping of plant gene expression and bacterial colonization has great potential in elucidating interactions between plant cells and bacterial cells at the single-cell level (Figure 7G). However, it is challenging or impossible to fully capture such interactions on a 2D tissue section, and thus a 3D analysis of both plant genes and bacterial colonization is critical. Recently, we developed a technology called PHYTOMap (Nobori et al., 2022), which can spatially map dozens of genes in 3D in whole-mount plant tissues. Such a method, combined with our spatiotemporal atlas, will open a new avenue toward a comprehensive characterization of cell populations identified in this study.

## Acknowledgments

We thank Kenichi Tsuda, James Walker, and Natanella Illouz-Eliaz for the critical comments on the manuscript, and Daniel Kliebenstein and Jeffrey Dangl for useful discussions. T.N. was supported by Human Frontiers Science Program (HFSP) Longterm Fellowship (LT000661/2020-L). J.R.E. is an Investigator of the Howard Hughes Medical Institute.

## Author contributions

T.N. conceived and designed the study and experiments with guidance from J.R.E. T.N., T.A.L, and J.N. performed experiments. T.N., A.M., and J.Z. analyzed data, T.N. wrote the initial draft of the manuscript, and A.M., T.A.L., J.Z., and J.R.E edited the manuscript.

## Data and code availability

The single-cell sequencing data used in this study are deposited in the National Center for Biotechnology Information Gene Expression Omnibus database (accession no. GSE226826). The code to analyze snMultiome and MERFISH data is available at https://github.com/tnobori/snMultiome and https://github.com/amonell/PlantMERFISH.

## Supplementary figures

**Figure S1.**
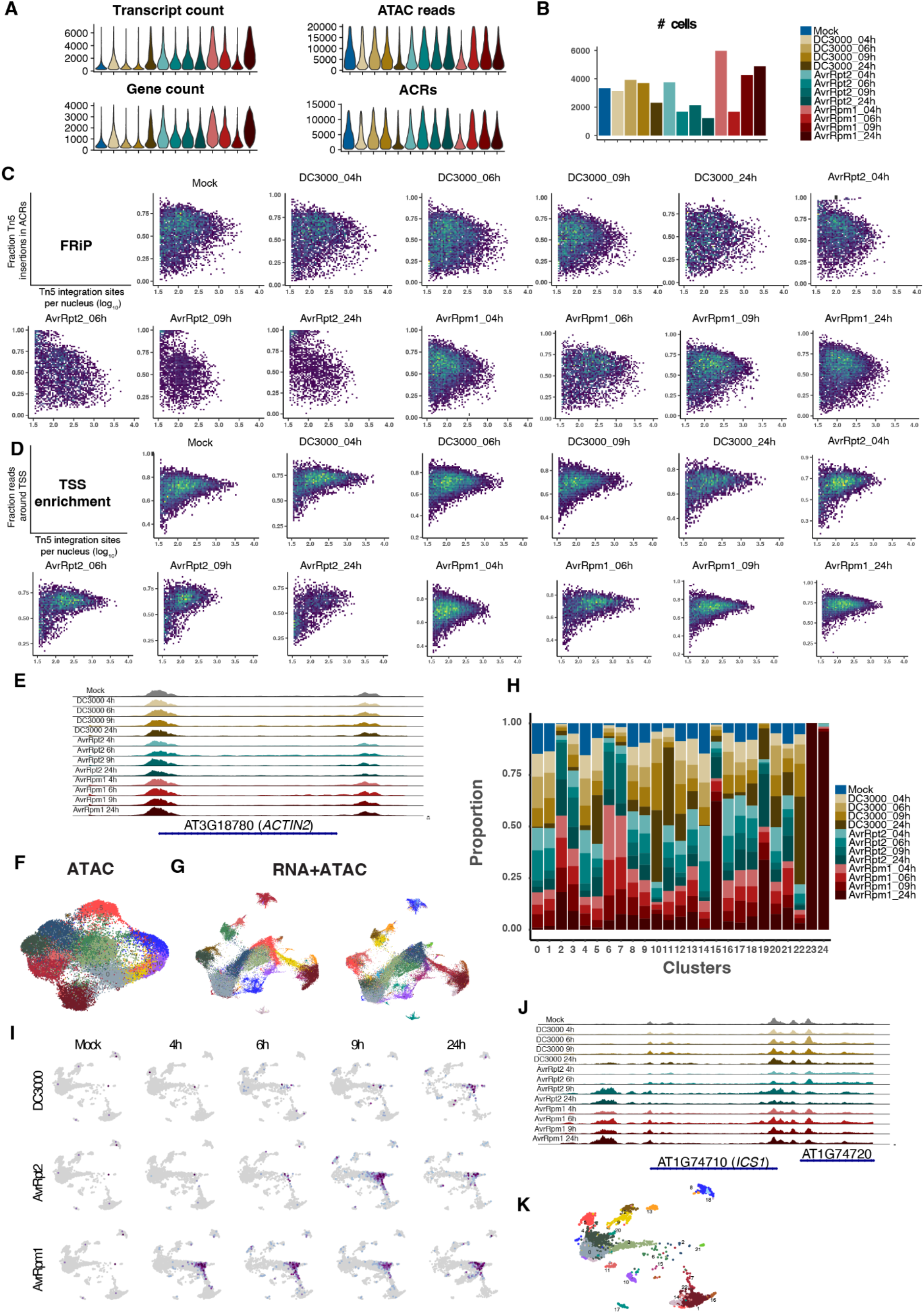
Quality control of snMultiome data, related to Figure 1. (A) Violin plots showing transcripts per nucleus, genes per nucleus, ATAC reads per nucleus, and accessible chromatin regions (ACRs) per nucleus in each sample. (B) The number of cells per sample. (C) Density scatterplots showing fraction reads in peaks (FRiP) score in each sample. x-axis: log10 transformed read depths. y-axis: fraction of Tn5 integration sites in ACRs. (D) Density scatterplots of log10 transformed read depths (x-axis) by the fraction of Tn5 integration sites mapping to within 2 kb upstream and 1 kb downstream of transcription start sites (TSSs) (y-axis). (E) Sample-aggregated chromatin accessibility around *ACTIN2*. (F) Two-dimensional embedding of chromatin accessibility similarity among nuclei from all samples with uniform manifold approximation and projection (UMAP). Nuclei are colored by Leiden clusters. (G) UMAP embeddings based on a joint neighbor graph that represents both gene expression and chromatin accessibility measurements. Nuclei are colored by *de novo* Leiden clusters based on the joint analysis (left) and Leiden clusters defined by gene expression measurement alone (right; Figure 1B). (H) Stacked bar plots showing the representation of gene expression-based Leiden clusters in each sample. (I) Expression of *ICS1* in each nucleus in each sample. (J) Sample-aggregated chromatin accessibility around *ICS1*. (K) UMAP plot showing cells in the mock sample.

**Figure S2.**
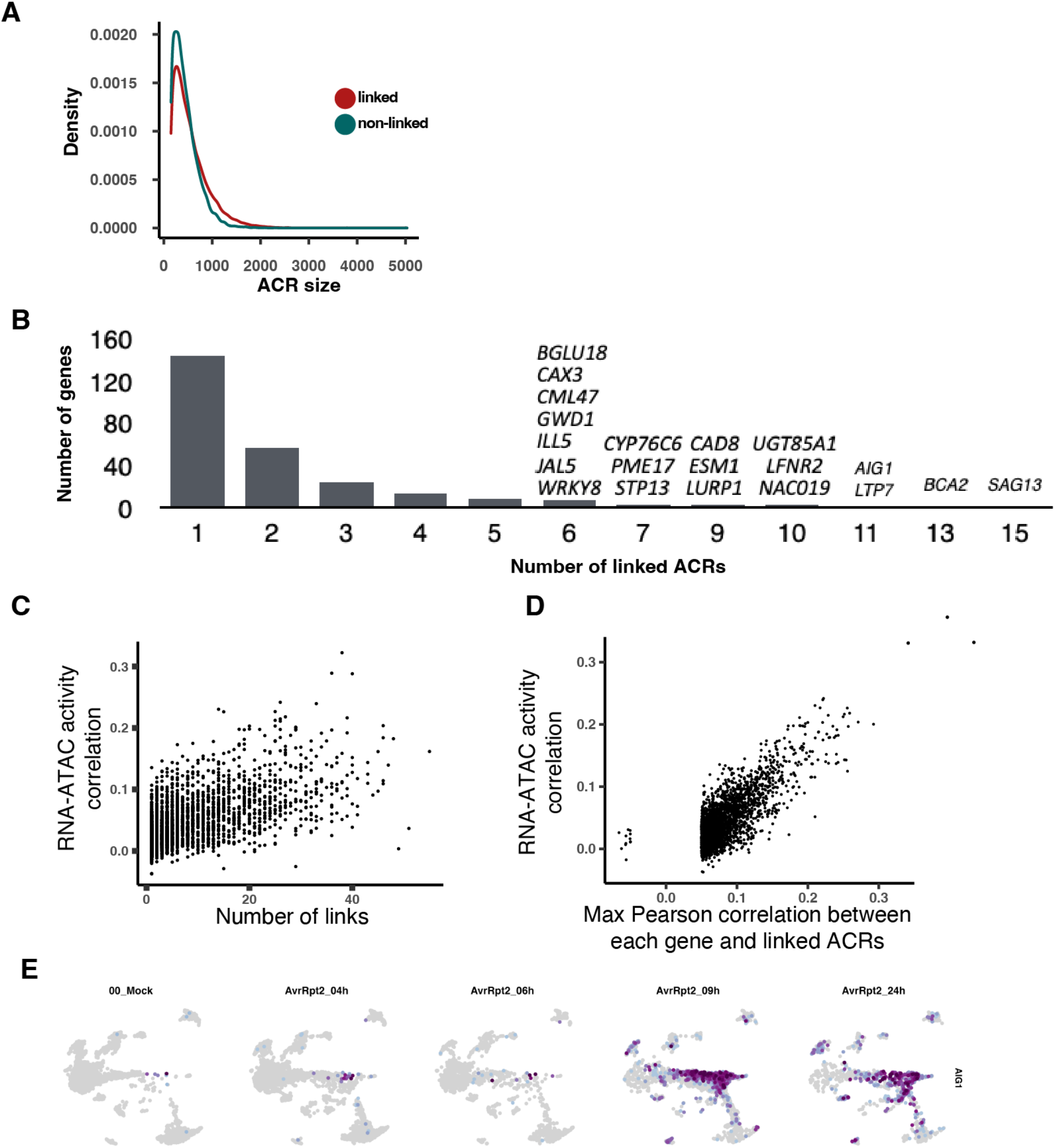
Links between transcriptome and chromatin accessibility, related to Figure 2. (A) Density plot showing the frequency of the size of ACRs for linked or non-linked genes. (B) Bar plot showing the number of genes with different numbers of linked ACRs. A full list is available in Table S3. (C) Scatter plot showing the relationship between the number of linkages (x-axis) and RNA-ATAC activity correlation score (y-axis; Figure 1I). (D) Scatter plot showing the relationship between the maximum correlation coefficient between each gene and linked ACRs (x-axis) and RNA-ATAC activity correlation score (y-axis; Figure 1I). (E) Expression of *AIG1* in plants infected by AvrRpt2.

**Figure S3.**
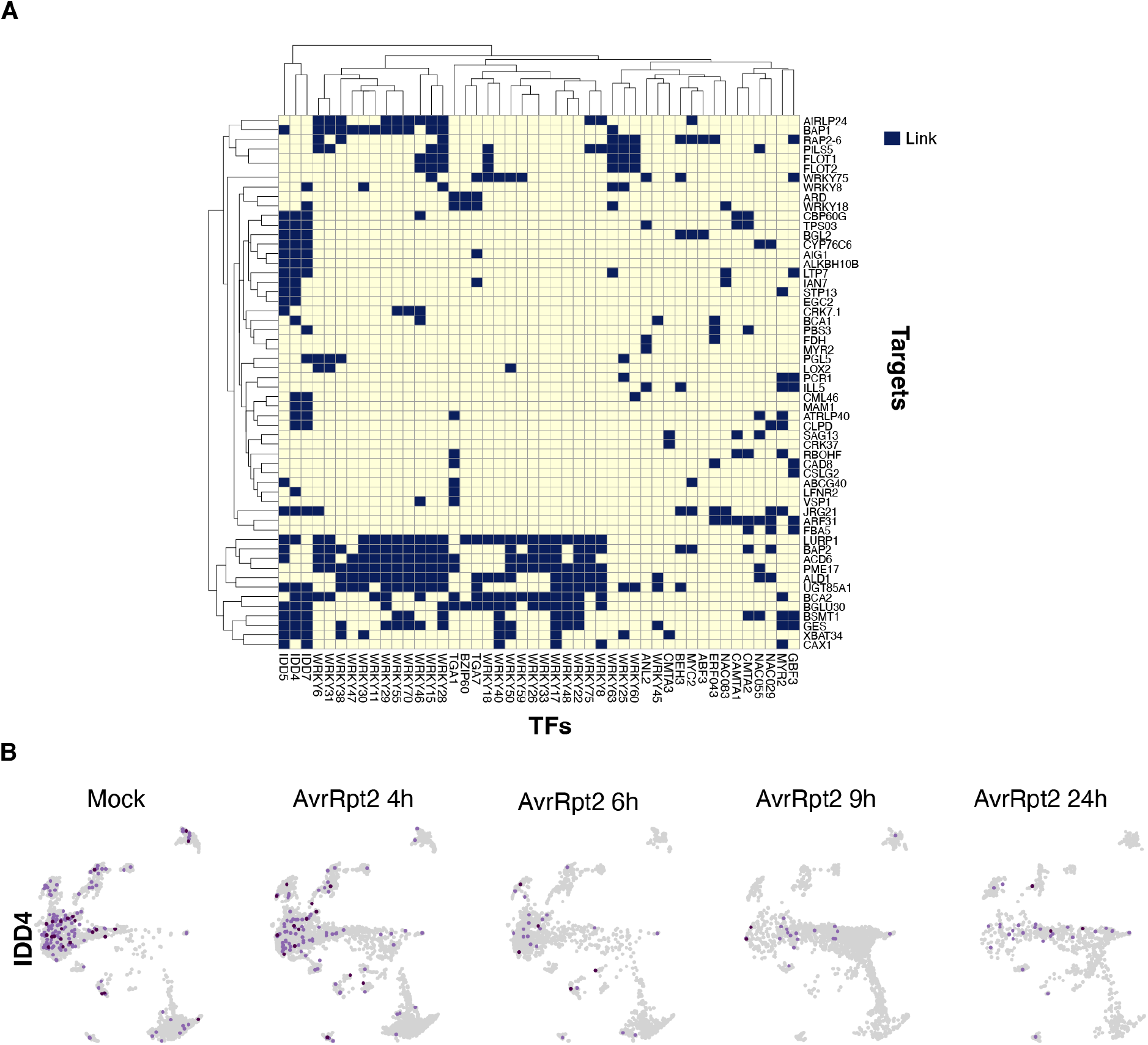
Identification of TF-gene modules, related to Figure 3. (A) Heatmap showing links between TFs and their predicted targets. (B) Expression of *IDD4* in plants infected by AvrRpt2.

**Figure S4.**
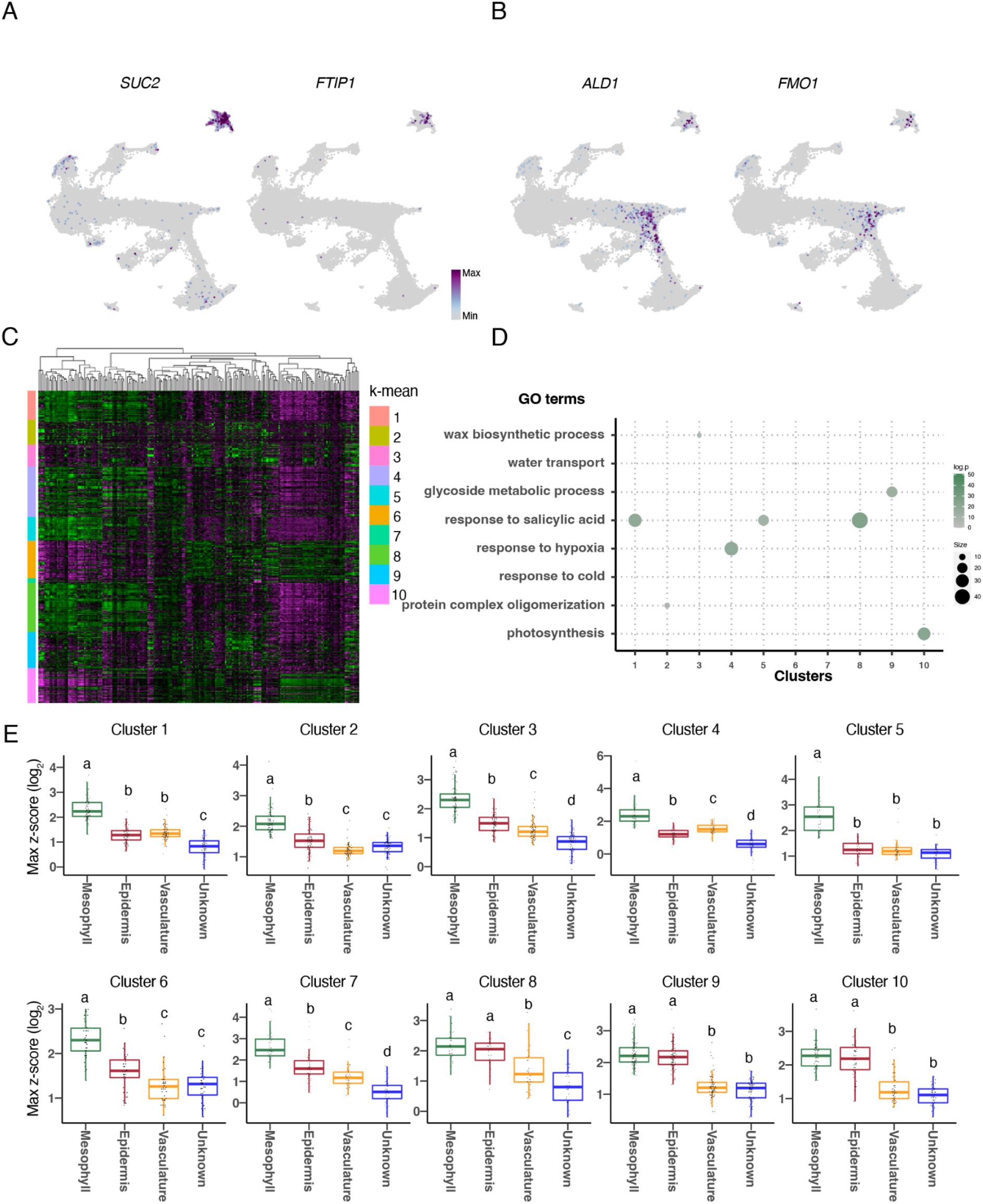
Analysis of sub-cell populations, related to Figure 4. (A) Expression of the phloem companion cell markers *SUC2* and *FTIP1*. (B) Expression of *ALD1* and *FMO1*. (C) Heatmap showing relative expression of genes in sub-clusters. The sidebar shows k-mean clusters. (D) GO enrichment analysis of genes in each k-mean cluster in (A). Genes in clusters (1, 5, and 8) were used for Figure 4F. (E) Boxplots showing the maximum z-score of genes belonging to each k-mean cluster (shown in C and Figure 4F) for each developmental cell type. Different letters indicate statistically significant differences (adjusted P < 0.05) based on a one-way ANOVA followed by Tukey’s HSD post hoc test.

**Figure S5.**
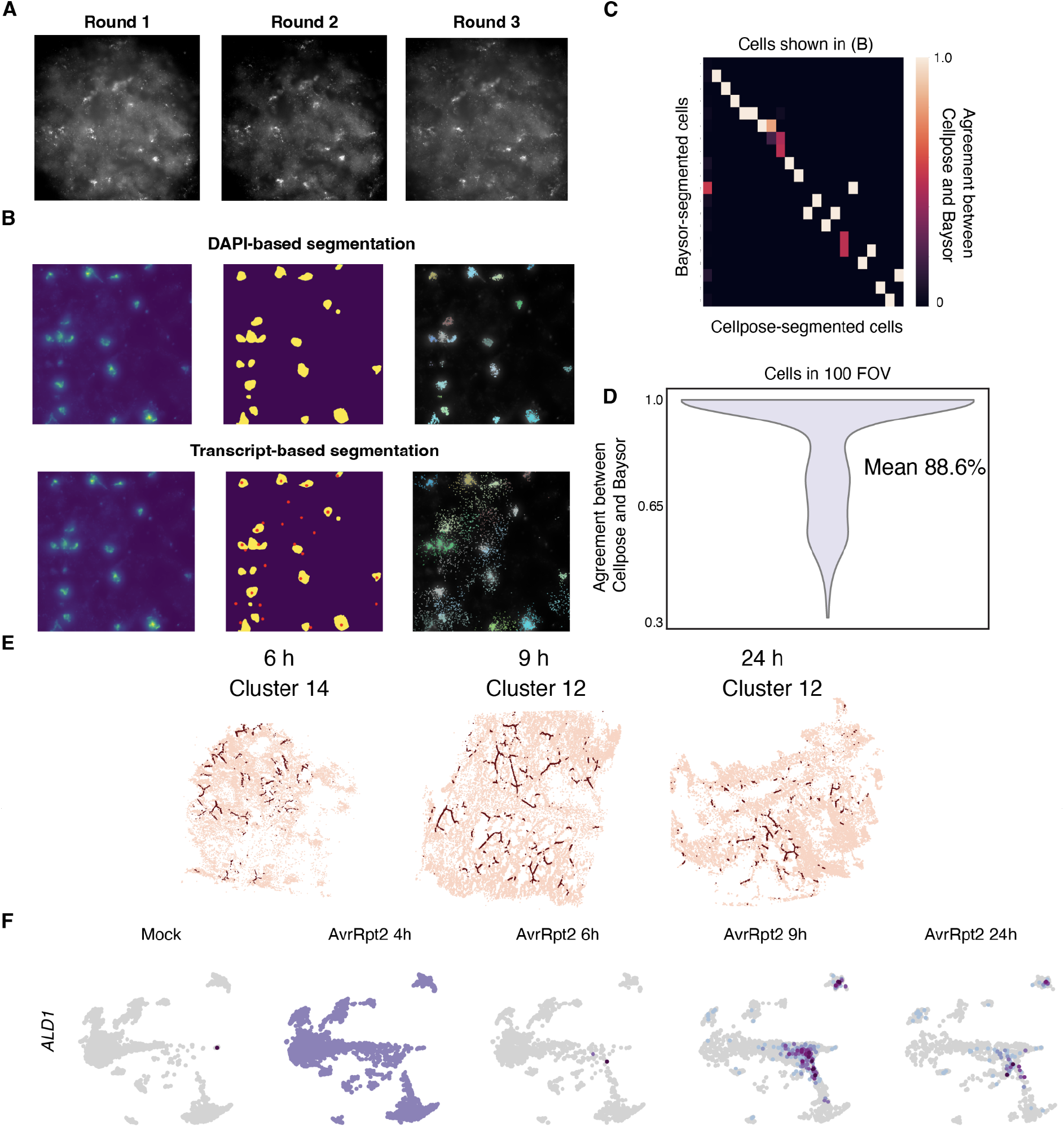
MERFISH data analysis, related to Figure 5. (A) Example raw images of MERFISH that capture the same region of tissue across three imaging rounds. White spots are signals derived from single mRNA molecules. (B) DAPI (top) and transcript (bottom) based segmentation. The left panels show DAPI images. In the middle panels, DAPI-based segmentation masks are shown in yellow. The centroids of cells detected by transcript-based segmentation are shown in red (in the lower bottom panel). The right panels show transcripts within segmentation masks colored differently for each cell. (C, D) The fraction of transcripts in Cellpose-segmented cells fell within Baysor-segmented cells. (C) Cells detected in the field of view (FOV) shown in (B) were used. (D) Cells detected in 100 FOV were used. (E) Spatial mapping of *de novo* MERFISH clusters annotated as vasculature. (F) Expression of *ALD1* in plants infected by AvrRpt2.

**Figure S6.**
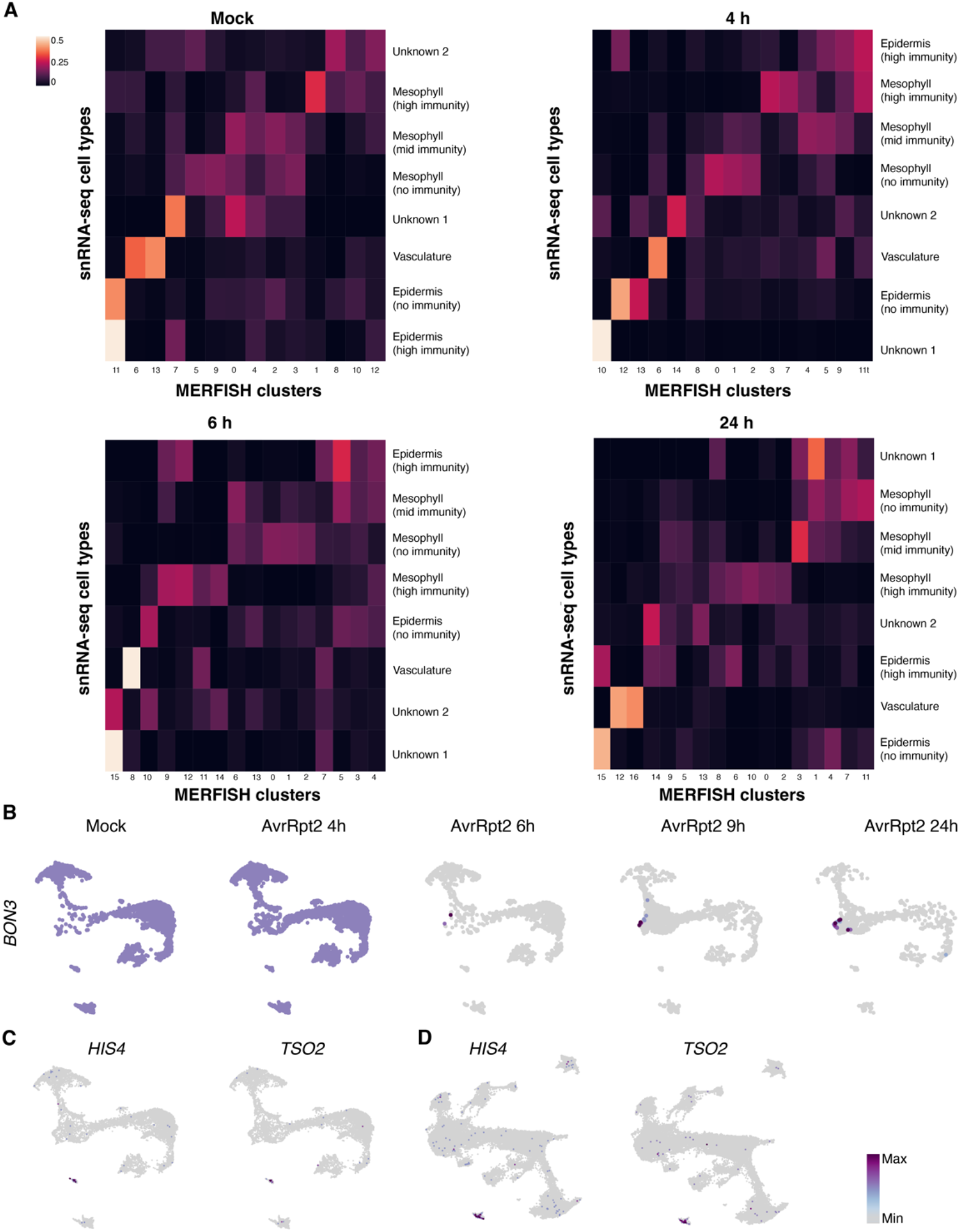
Integration between MERFISH and snMultiome. (A) Proportion of *de novo* MERFISH clusters (Mock, 4 h, 6 h, and 24 h; shown in Figure 5E) assigned to cell populations defined by snRNA-seq based on data integration (see Method). (B) Expression of *BON3* in plants infected by AvrRpt2 shown in the UMAP in Figure 6B. (C, D) Expression of S phase markers, *HIS4* and *TSO2*, shown in the UMAP in (C) Figure 6B and (D) Figure 1B.

**Figure S7.**
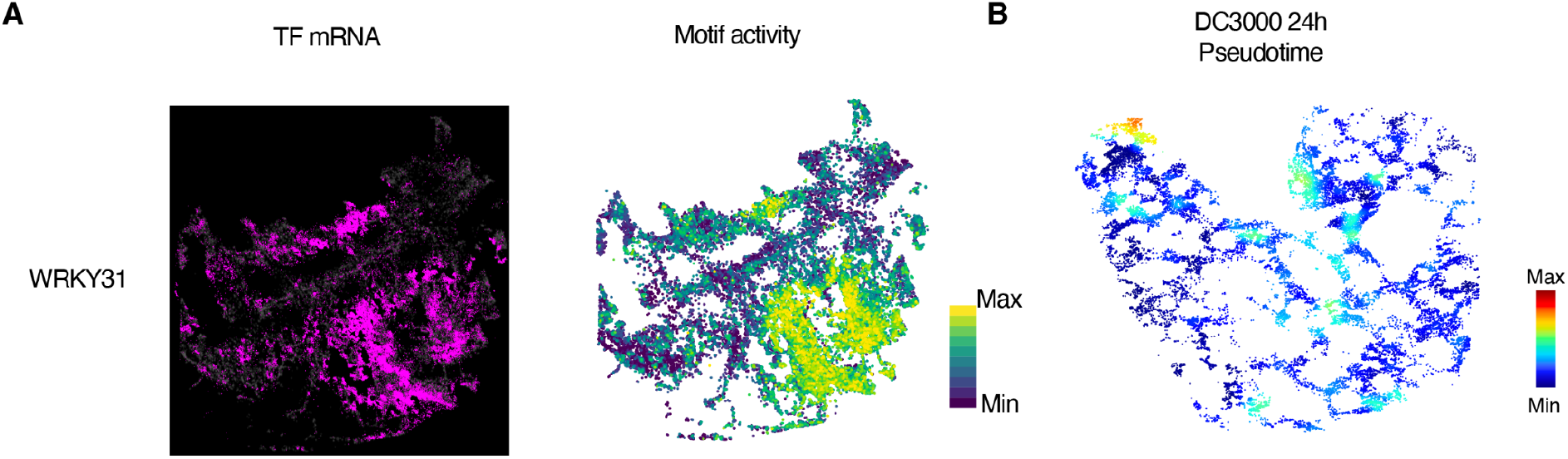
imputation of snMultiome data into MERFISH tissues, related to Figure 7. (A) (Left) Imputed motif enrichment scores of *WRKY31*. (Right) mRNA expression of *WRKY31* by MERFISH. (B) Spatial mapping of pseudotime values based on data integration and label transfer between snRNA-seq and MERFISH data of DC3000- infected plants.

## Supplementary tables

Table S1: *De novo* marker gene of individual clusters, related to Figure 1

Table S2: Significant links between accessible chromatin regions and genes, related to Figure 2

Table S3: Number of links per gene, related to Figure 2

Table S4: List of linked defense genes and non-linked defense genes, related to Figure 2

Table S5: Cluster specific motifs, related to Figure 3

Table S6: List of MERFISH target genes, related to Figure 5

## Methods

### Reagents and kits

Ammonium persulfate (Sigma, 09913), Bovine Serum Albumin (BSA) solution (Sigma, A1595), Chromium Next GEM Single Cell Multiome ATAC + Gene Expression (10X Genomics, PN-1000283), Corning™ Falcon™ Cell Strainers (Corning, 08-771-2) DTT (dithiothreitol) (Thermo, R0861), EDTA, pH 8.0 RNase -free (Invitrogen, AM9260G), KCl (2 M) RNase-free (Invitrogen, Invitrogen), MACS® SmartStrainers (Milteny Biotec, 130-098-458), MERSCOPE 500 Gene Imaging Kit (Vizgen, 10400006), MERSCOPE 500 Gene Panel (Vizgen, 10400003), MERSCOPE Sample Prep Kit (Vizgen, 10400012), N,N,N’,N’-Tetramethylethylenediamine (TEMED) (Sigma, T7024-25ML), NaCl (5 M) RNase-free (Invitrogen, AM9759), NP40 (IGEPAL® CA-630) (Sigma, I8896), Paraformaldehyde (Sigma, F8775), Phosphate-Buffered Saline (10X) pH 7.4 RNase-free (Thermo Fisher, AM9625), Protease Inhibitor Cocktail (Sigma, P9599), Protector RNAse inhibitor (Sigma, 3335402001), Scigen Tissue-Plus™ O.C.T. Compound (Fisher, 23-730- 571), Spermidine (Sigma, S2626), Spermine (Sigma, 85590), Triton-X (Sigma, 93443), UltraPure™ 1 M Tris-HCI Buffer pH 7.5 (Invitrogen, 15567027)

### Plant growth and bacterial infection

*Arabidopsis thaliana* Col-0 was grown in a chamber at 22°C with a 12-h light period and 60-70% relative humidity for 30-31 days. Bacterial strains were cultured in the King’s B liquid medium with antibiotics (Rifampicin and Tetracyclin) at 28°C. Three bacterial strains–*Pseudomonas syringae* pv. *tomato* DC3000 carrying the empty vector (pLAFR3), avrRpt2 (pLAFR3), and avrRpm1 (pLAFR3)–were described previously (Whalen et al., 1991; Kunkel et al., 1993; Debener et al., 1991). Bacteria were harvested by centrifugation and resuspended in sterile water to an OD_600_ of 0.001 (approximately 5 x 10^5^ CFU ml^-1^). In total, 20 *A. thaliana* leaves (four fully-expanded leaves per plant) were syringe-inoculated with bacterial suspensions using a needleless syringe. For each strain, four-time points (4, 6, 9, and 24 h) were sampled at the same time of the day by infiltrating bacteria at different times. The 20 infected leaves were harvested with forceps and immediately processed for nuclei extraction. For the mock condition, water-infiltrated leaves were harvested after 9 h.

### Nuclei extraction and single-nucleus sequencing

Fresh Nuclei purification buffer (NPB; 15 mM Tris pH 7.5, 2 mM EDTA, 80 mM KCl, 20 mM NaCl, 0.5 mM spermidine, 0.2 mM spermine, 1:100 BSA, and 1:100 protease inhibitor cocktail) was prepared before the experiment and chilled on ice. All the following procedures were performed on ice or at 4°C. Twenty leaves were chopped in 500-1,000 µL of cold NPB with 1:500 Protector RNase IN with a razor blade on ice for 5 min to release nuclei and incubated in 20 mL of NPB. The crude nuclei extract was sequentially filtered through 70 µm and 30 µm cell strainers (70 µm: Corning™ Falcon™ Cell Strainers, Corning, 08-771-2; 30 µm: MACS® SmartStrainers, Milteny Biotec, 130-098-458). Triton-X and NP40 were added to the extract at the final concentration of 0.1% each, and the extract was incubated at 4°C for 5 min with rotation. The nuclei suspension was centrifuged at 50 x g for 3 min in a swing-rotor centrifuge to pellet non-nuclei debris, and the supernatant was taken. Nuclei were pelleted by centrifugation at 500 x g for 5 min with a swing-rotor centrifuge. When the pellet was green, the pellet was resuspended in 20 mL of NPBd (NPB, 0.1% Triton-X, and 0.1% NP40) with 1:1000 Protector RNase IN by pipetting, followed by centrifugation at 500 x g for 5 min. When the pellet was translucent, the NPBd wash was skipped. The pellet was then washed by resuspending it in 20 mL of NPB with 1:1000 Protector RNase IN and centrifuging at 500 x g for 5 min with a swing-rotor centrifuge. The pellet was resuspended in 950 µL of 1X Nuclei Buffer (10X Genomics; PN-2000207) with 1:40 Protector RNase IN and 1mM DTT. The nuclei suspension was centrifuged at 50 x g for 3 min in a swing-rotor centrifuge to pellet non-nuclei debris, and the supernatant was taken. This step was repeated one more time. Then, nuclei were counted manually with a hemocytometer. Nuclei were pelleted by centrifugation at 500 x g for 5 min with a swing-rotor centrifuge and removed supernatant leaving approximately 10 µL of the buffer. Nuclei were counted again, and up to 16k nuclei were used for the following steps. Single-cell RNA-seq and ATAC-seq libraries were constructed according to the manufacturer’s instructions (10X Genomics, catalog: CG000338). Single-nucleus RNA-seq libraries were sequenced with Illumina NovaSeq 6000 in dual-index mode with 10 cycles for i7 and i5 index.

Single-nucleus ATAC-seq libraries were also sequenced with Illumina NovaSeq 6000 in dual-index mode with eight and 24 cycles for i7 and i5 index, respectively.

## Single-cell multiomics analysis

### Raw data processing

Sequence data were processed to obtain single-cell feature counts by running *cellranger* (v6.0.1) and *cellranger-arc* (v2.0.0) for snRNA-seq and snATAC-seq, respectively. For snRNA-seq, the –include-introns option was used to align reads to the *A. thaliana* nuclear transcriptome built with the TAIR10 genome and Araport 11 transcriptome. Count data were analyzed with the R packages *Seurat v4* (Hao et al., 2021) and *Signac* (Stuart et al., 2021).

### Quality control and cell filtering

QC matrices for snATAC-seq were generated by a modified version of *loadBEDandGenomeData* function in the R package *Socrates* (Marand et al., 2021). ACRs were identified with MACS2 (Zhang et al., 2008) with the following parameters: -g (genomesize)=0.8e8, shift=-50, extsize=100, and --qvalue=0.05, --nomodel, --keep-dup all. Fraction of reads mapping to within 2 kb upstream or 1 kb downstream TSS was calculated. Nuclei were filtered with the following criteria: 200< RNA UMI count <7000, RNA gene count >180, 200< ATAC UMI count <20000, fraction of RNA reads mapped to mitochondrial genome <5%. Seurat objects of individual samples were merged with the *Merge_Seurat_List* function of the *scCustomize* package.

### snRNA-seq clustering

snRNA-seq clustering was performed with the R package *Seurat*. Cell-by-gene RNA count matrix was normalized by *SCTransform*. Dimension reduction was performed by principal component analysis (PCA) with *RunPCA*. Technical variance among samples was reduced with *Harmony* (Korsunsky et al., 2019) using PCs 1 to 20. Graph-based clustering was performed on the Harmony corrected PCs 1 to 20 by first computing a shared nearest-neighbor graph using the PC low-dimensional space (with k=20 neighbors) and then applying Louvain clustering (resolution=1.0), and projected into an additionally reduced space with UMAP (n.neighbors=30L, and min.dist=0.01).

### snATAC-seq peak calling

Peaks were called independently on each cluster defined by snRNA-seq data and then combined using the *CallPeaks* function of *Signac*, which uses MACS2 with the following parameters: effective.genome.size=1.35e8, extsize=15, shift=-75. Peak counts were quantified with the *FeatureMatrix* function.

### snATAC-seq clustering

Dimensionality reduction was performed by latent semantic indexing (LSI) (Cusanovich et al., 2015). First, the top 95% most common features were selected with the *FindTopFeatures* function. Then, the term-frequency inverse-document-frequency (TF-IDF) was computed with *RunTFIDF* with scale.factor=100000. The resulting TF-IDF matrix was decomposed with singular value decomposition with *RunSVD*, which uses the *irlba* R package. Technical variance among samples was reduced with *Harmony* using LSI components 2 to 10. Graph-based clustering was performed on the Harmony-corrected LSI components 2 to 10 by first computing a shared nearest-neighbor graph using the LSI low-dimensional space (with k=20 neighbors) and then applying Louvain clustering (resolution=0.8), and projected into an additionally reduced space with UMAP (n.neighbors=30L, and min.dist=0.01).

### RNA-ATAC joint clustering

The two modalities were integrated with weighted nearest neighbor analysis with *FindMultiModalNeighbors* of *Seurat* using Harmony-corrected PC 1 to 20 for RNA and Harmony-corrected LSI components 2 to 10 for ATAC. Then, SLM (resolution = 0.5) was applied and projected with UMAP (n.neighbors=30L, and min.dist=0.1).

### ATAC gene activity score

ATAC gene activity score was calculated with the *GeneActivity* function of *Signac* with extend.upstream=400.

### Linkage analysis

*LinkPeaks* of *Signac* was used to call significant peak-to-gene linkage for each infection condition (mock, DC3000, AvrRpt2, and AvrRpm1). Background corrected Pearson correlation coefficient between the gene expression of each gene and the accessibility of each peak within 500 kb of the gene TSS was calculated. A *P* value was calculated for each peak-gene pair using a one-sided z-test, and peak-gene pairs with a *P* value <0.05 and a Pearson correlation coefficient >0.05 was retained as a significant link.

### Motif enrichment analysis

Motifs present in the JASPAR2020 database (Fornes et al., 2020) for Arabidopsis (species code 3702) were used. Cluster-specific peaks were first identified with *FindMarkers* of *Seurat* with default parameters. A hypergeometric test was used to test for the overrepresentation of motifs in the set of differentially accessible peaks using *FindMotifs* of *Signac*. Motif plots were generated with *MotifPlot*. Motif enrichment scores (motif deviation scores) of individual TF motifs in individual cells were calculated with *chromVAR* (Schep et al., 2017).

### Sub-cluster analysis

Nuclei with the same sub-cluster label were aggregated, and log_2_-transformed transcripts per million (TPM) was calculated. Sub-clusters with more than 20,000 undetected genes were removed from the analysis. For the sub-clusters that passed the filtering, genes were clustered by k-mean clustering with k=10 (Figure S4C). Then, GO enrichment analysis was performed for the genes in each cluster (Figure S4D). Three clusters (1, 5, and 8) showed the enrichment of an immunity-related function (responses to salicylic acid); genes in these clusters were defined as “putative immune genes” and used for the downstream analysis.

### Pseudotime analysis

To calculate pseudotime, the cell-by-gene matrix for cells in clusters {0, 4, 6, 8, 9, 11}, which were considered to be mesophyll cells, was obtained from the snMultiome data. Using the Python pseudotime package *Palantir*, PCA was calculated using *palantir.utils.run_pca* with use_hvg=False, and n_components=10. Diffusion maps and multiscale space were determined using *palantir.utils.run_diffusion_maps* with n_components=5, and *palantir.utils.determine_multiscale_space*, respectively. To denoise the gene expression matrix, we used *palantir.utils.run_magic_imputation* for MAGIC imputation. Looking at the multiome UMAP (Figure 6B), we set the rightmost Mock cell, belonging to cluster 0, as the Palantir starting cell. The leftmost 24 hpi cell, belonging to cluster 11, and the uppermost 24 hr cell, belonging to cluster 6, were set as terminal states. *Palantir.core.run_palantir* was initiated with these starting and terminal states, along with the parameters knn=20, and num_waypoints=100. Heatmap gene trends were calculated with *palantir.preresults.compute* gene trends.

## MERFISH

### MERFISH panel design

We curated 500 target genes, including (1) *A. thaliana* leaf cell type markers defined by (Procko et al., 2022), (2) genes involved in various processes such as immunity, hormone pathways, and epigenetic regulation, and (3) a variety of TFs previously analyzed with DAP-seq (a TF-DNA interaction assay) (O’Malley et al., 2016). Genes for which more than 25 specific probes could not be designed based on Vizgen’s probe design software were excluded from the target gene panel. Highly expressed genes could cause the overcrowding of smFISH signal and hinder MERFISH quantification. To avoid including highly expressed genes in the panel, we assessed target gene expression by using a publicly available bulk RNA-seq dataset of *A. thaliana* infected by AvrRpt2 in the same setup as the present study at eight different time points (1, 3, 4, 6, 9, 12, 16, and 24h) (Mine et al., 2018). For each gene, the highest expression value among the eight time points was used. Genes that showed the transcript per million (TPM) >710 were not included in the panel. The total TPM of the 500 genes was approximately 22,000. *ICS1* was targeted with a single round smFISH as the gene is highly expressed. Bacterial cells were visualized by targeting 19 highly expressed genes (based on *in planta* bulk RNA-seq data by (Nobori et al., 2018b)) as a single target. Table S6 shows a list of genes targeted by MERFISH. All the probes were designed and constructed by Vizgen.

### Tissue sectioning, fixation, and mounting

Plants were grown according to the previously described methods. Leaves matching the aforementioned treatments and timepoints were excised and immediately incubated and acclimated in OCT (Fisher) for 5 min. Following incubation, the leaves were immediately frozen according to Giacomello and Lundeberg, 2018. Tissue blocks were acclimated to −18°C in a pre-cooled cryostat chamber (Leica) for 1 h. Tissue blocks were trimmed until the tissue was entered, after which 10 µm sections were visually inspected until the region of interest was exposed. Sample mounting and preparation were performed according to the MERSCOPE user guide, with slight modifications. Briefly, a 10 µm section was melted and mounted onto a room temperature MERSCOPE slide (Vizgen, 20400001), placed into a 60 mm Petri dish and re-frozen by incubation in the cryostat chamber for 5 min. Subsequent steps were performed with the mounted samples in the Petri dish. The samples were then baked at 37°C for 5 min and were then incubated in fixation buffer (1X PBS, 4% formaldehyde) for 15 min at RT. Samples were then washed with 1X PBS containing 1:500 RNAse inhibitor (Protector RNAse inhibitor, Millipore Sigma) for 5 min at RT in triplicate. Following the final PBS wash, samples were dehydrated by incubation in 70% EtOH at 4°C overnight.

### MERFISH experiment

Tissue sections were processed following Vizgen’s protocol. After removing 70% ethanol, the sample was incubated in the Sample Prep Wash Buffer (PN20300001) for 1 min then incubated in the Formamide Wash Buffer (PN20300002) at 37°C for 30 min. After removing the Formamide Wash Buffer, the sample was incubated in MERSCOPE Gene Panel Mix at 37°C for 42 h. After the probe hybridization, the sample was washed twice with the Formamide Wash Buffer at 47°C for 30 min and once with the Sample Prep Wash Buffer at RT for 2 min. After the washing, the sample was embedded in hydrogel by incubating in the Gel Embedding Solution (Gel Embedding Premix (PN20300004), 10% w/v ammonium persulfate solution, and N,N,N’,N’-tetramethylethylenediamine) at RT for 1.5 h. Then, the sample was cleared by first incubating in the Digestion Mix (Digestion Premix (PN 20300005) and 1:40 Protector RNase inhibitor) at RT for 2 h, followed by the incubation in the Clearing Solution (Clearing Premix (PN 20300003) and Proteinase K) at 47°C for 24 h then at 37°C for 24 h. The cleared sample was washed twice with the Sample Prep Wash Buffer and stained with DAPI and PolyT Staining Reagent at RT for 15 min then washed with the Formamide Wash Buffer at RT for 10 min and rinsed with the Sample Prep Wash Buffer. The sample was imaged with the MERSCOPE Instrument, and detected transcripts were decoded on the MERSCOPE Instrument using a Codebook generated by Vizgen. Transcripts were visualized on Vizgen’s Visualizer.

### MERFISH segmentation and processing

Cell boundary segmentation was performed for each MERSCOPE data output. DAPI and poly(A)-targeting probes demonstrated variable success in staining nuclei and cytoplasm, respectively, depending on the samples and tissue regions examined (Figure 5C and Figure S5B show failed and successful segmentation, respectively). In contrast, dense transcript areas marked nuclei locations more robustly (Figure S5B). Therefore, a transcript-based segmentation was used. For each sample, a 2D Numpy array of “0”s was generated, modeling the total pixel area imaged. The coordinates of identified RNA transcripts were changed from “0” to “1” within this array. Next, the array was blurred using OpenCV’s *cv2.GaussianBlur function* with ksize=(5,5). The resulting array was chunked into 2000×2000 pixel regions. These regions were loaded into *Cellpose* (Pachitariu and Stringer, 2022), a deep learning-based segmentation tool, and a custom segmentation model was trained by manually segmenting nuclei objects across ten 2000×2000 pixel regions in the Cellpose GUI.

The custom model was then used to predict the segmentation boundaries within all the remaining regions, with the parameters diameter=22.92, flow_threshold=0.7, and cell probability threshold=-2. Next, the total number of cells in an experiment was calculated by summing the number of unique cells across all regions. This number was then used to initialize the --num-cells-init command in another segmentation tool, *Baysor* (Petukhov et al., 2022), which considers the joint likelihood of transcriptional composition and cell morphology to predict cell boundaries. Baysor was run using a downloaded Docker image and parameters -s 250, --n clusters 1, -i 1, --force-2d, min-molecules-per-gene=1, min-molecules-per-cell=50, scale=250, scale-std=“25%”, estimate-scale-from-centers=true, min-molecules-per-segment=15, new-component-weight=0.2, new-component-fraction=0.3.

To test the quality of our transcript-based segmentation, we used a FOV with successful DAPI staining (which was rare in our samples) and performed a DAPI-based watershed segmentation and transcript-based segmentation (Figure S5B). Results from these two segmentation strategies agreed with each other in general, with the transcript-based approach could capture transcripts in the cytoplasm in addition to those in the nucleus (Figure S5B-D), indicating that our segmentation approach can reliably capture cells.

After *Baysor* segmentation, a cell-by-gene matrix was created from the transcript cell assignments. Cells with fewer than 50 assigned transcripts were removed. *Scanpy* was used for post-processing of our MERFISH experiments. After loading the respective cell by gene matrix into an Anndata object for each experiment, we stored the spatial coordinates of each cell obtained from *Baysor*. The individual transcript counts in each cell was normalized by the total number of transcript counts per cell. The Anndata cell by gene matrix was then log-scaled.

### MERFISH-snMultiome data integration and label transfer

To integrate the MERFISH experiment data with our snMultiome data, we first concatenated all MERFISH experiments into a single Anndata object, normalized the total read counts, and log-transformed the data. We selected the 500 genes probed in the MERFISH experiments from the RNA data of snMultiome cells (X_rna_) to fit a PCA model with 50 components and then projected both the RNA data and MERFISH data (X_merfish_) to the embedding space with this PCA model to generate PC_rna_ and PC_merfish_. We used ALLCools (https://github.com/lhqing/ALLCools) to perform the integration, which has a python version of Seurat v3 that is more scalable than the original R version. Specifically, we used PCA with 30 components to find mutual 5 nearest neighbors as anchors between X_rna_ and X_merfish_, and project the MERFISH embedding (PC_merfish_) to the RNA embedding (PC_rna_) to generate corrected MERFISH embedding with the anchors. The anchors were also used to transfer the cell type labels, whole transcriptome, and motif accessibility of snMultiome cells to the MERFISH cells, to visualize the spatial distribution of cell types, non-probed genes, and epigenome signatures. Specifically, for each MERFISH cell as a query, we found its 100 nearest anchors, and the data from the 100 snMultiome cells on these anchors were averaged and then weighted by the distance between the MERFISH query cell and the MERFISH cell on the corresponding anchor. For continuous data, the values of the 100 snMultiome cells on the nearest anchors were weighted averaged. For categorical data, we used one-hot encoding to represent the data, and the data vectors corresponding to the 100 snMultiome cells on the nearest anchors were weighted averaged. The resulting vector represents the probability of the query cell belonging to each category, and the category with the maximum probability is used as the final assignment. To plot the joint UMAP embedding of both modalities (MERFISH and snRNA-seq), we first used the *Harmony* function *run_harmony*, with nclust = 50 and vars_use = [‘modality’]. We then used the function *sc.tl.umap* to derive a UMAP from the Harmony PCA coordinates.

### Comparison between MERFISH clusters and snRNA-seq clusters (Figure 6A related)

In order to generate Leiden clusters from the MERFISH data, we performed a *de novo* clustering of each experiment (Figure 5F). Additionally, we transferred multiome cluster labels from the data integration between the MERFISH and multiome experiments. This resulted in each cell from each MERFISH experiment being assigned both a *de novo* cluster assignment based on the MERFISH data and a transferred cluster assignment based on the multiome data. To create the heatmaps in Figure 6A, we show the percentage of cells belonging to each transferred cluster that simultaneously belong to a specific *de novo* cluster.

### smFISH quantification and bacterial colony identification

Quantification of transcripts labeled by smFISH was performed using the Python package *Big-FISH*. Seven z-planes of MERSCOPE smFISH images were projected into 2 dimensions by using *numpy.max* along the z-axis, and chunked into 2000×2000 regions. Spots were then called using the function *bigfish.detection.detect_spots* with threshold=50, spot_radius=(10, 10), and voxel_size=(3, 3). To identify bacterial colonies, *bigfish.detection.detect_spots* was called to label “bacterial meta gene” locations with threshold=200, log_kernel_size=(1.456, 1.456), and minimum_distance=(1.456, 1.456). To account for autofluorescence from the plant tissue, the same spot-caller was used to call spots on DAPI and *ICS1* channels. The spots from all three channels were aggregated, and *DBSCAN* from *scikit-learn.cluster* was used on all spots with eps=35 and min_samples=5. We kept all DBSCAN clusters where the DAPI and *ICS1* spots comprised less than 30% of the total cluster spots. These remaining clusters represented potential bacterial colonies. We manually evaluated each cluster to merge those marking the same colony and removed the clusters marking obvious autofluorescence e.g., signal from stomata. Windows of size 300×300 pixels were generated around each final cluster, and detect_spots was used with threshold=95, spot_radius=(17, 17), and voxel_size=(3, 3) for accurate quantification of individual bacteria per cluster.

### Bacterial neighborhood analysis (related to Figure 7G)

To determine the level of immunity of the neighborhood around each bacteria colony, we identified the five nearest cells in proximity to the colony. Then, the smoothed imputed pseudotime values of these neighboring cells were averaged to obtain a single mean pseudotime value. This value serves as an indicator of the overall immunity level of the area around each colony.

## Notes

### Competing Interest Statement

The authors have declared no competing interest.

### Summary of Updates

Figure 4 and References have been revised.

